# Haploid-resolved and chromosome-scale genome assembly in hexa-autoploid sweetpotato (*Ipomoea batatas* (L.) Lam)

**DOI:** 10.1101/2022.12.25.521700

**Authors:** Ung-Han Yoon, Qinghe Cao, Kenta Shirasawa, Hong Zhai, Tae-Ho Lee, Masaru Tanaka, Hideki Hirakawa, Jang-Ho Hahn, Xiangfeng Wang, Ho Soo Kim, Hiroaki Tabuchi, An Zhang, Tae-Ho Kim, Hideki Nagasaki, Shizhuo Xiao, Yoshihiro Okada, Jae Cheol Jeong, Soichiro Nagano, Younhee Shin, Hyeong-Un Lee, Sul-U Park, Seung Jae Lee, Keunpyo Lee, Jung-Wook Yang, Byoung Ohg Ahn, Daifu Ma, Yasuhiro Takahata, Sang-Soo Kwak, Qingchang Liu, Sachiko Isobe

## Abstract

Sweetpotato (*Ipomoea batatas* (L.) Lam) is the world’s seventh most important food crop by production quantity. Cultivated sweetpotato is a hexaploid (2n = 6x = 90), and its genome (B1B1B2B2B2B2) is quite complex due to polyploidy, self-incompatibility, and high heterozygosity. Here we established a haploid-resolved and chromosome-scale *de novo* assembly of autohexaploid sweetpotato genome sequences. Before constructing the genome, we created chromosome-scale genome sequences in *I. trifida* using a highly homozygous accession, Mx23Hm, with PacBio RSII and Hi-C reads. Haploid-resolved genome assembly was performed for a sweetpotato cultivar, Xushu18 by hybrid assembly with Illumina paired-end (PE) and mate-pair (MP) reads, 10X genomics reads, and PacBio RSII reads. Then, 90 chromosome-scale pseudomolecules were generated by aligning the scaffolds onto a sweetpotato linkage map. *De novo* assemblies were also performed for chloroplast and mitochondrial genomes in *I. trifida* and sweetpotato. In total, 34,386 and 175,633 genes were identified on the assembled nucleic genomes of *I. trifida* and sweetpotato, respectively. Functional gene annotation and RNA-Seq analysis revealed locations of starch, anthocyanin, and carotenoid pathway genes on the sweetpotato genome. This is the first report of chromosome-scale *de novo* assembly of the sweetpotato genome. The results are expected to contribute to genomic and genetic analyses of sweetpotato.

## Introduction

Sweetpotato (*Ipomoea batatas* (L.) Lam) is the seventh most important food crop in the world by production quantity^1^. Total global production of sweetpotato in 2020 was 89.5 million tons from 7.4 million ha. Sweetpotato has many advantages over other starch crops in terms of yield, nutritional value, and environmental adaptability to marginal lands, including desertification areas. Thus, sweetpotato is widely cultivated in areas ranging from the tropical to temperate zones for human consumption, animal feed, and industrial purposes. At high latitude, sweetpotato plants require less treatment with chemical pesticides and fertilizers. The minimum requirement for sweetpotato cultivation is a frost-free period lasting at least 4 months. Recently sweetpotato has been re-evaluated as a valuable medicinal plant with antiaging, anticancer, antidiabetic, and anti-inflammatory properties, since it contains high levels of low molecular weight antioxidants such as vitamin C and carotenoids, dietary fiber and potassium^2^. In addition, sweetpotato has higher carbohydrate content than other starch crops, indicating high potential for bioethanol production on marginal lands^3^.

Sweetpotato originated in Central and South America and is one of the oldest domesticated crops in the Americas^4^. The cultivated sweetpotato and its closest wild relatives belong to the genus *Ipomoea* (family Convolvulaceae). The wild ancestors of sweetpotato have not been fully identified, but morphologic, cytogenetic, and molecular studies all support a close relationship with *I. trifida* (2n = 2x = 30)^5^. The cultivated sweetpotato is a hexaploidy (2n = 6x = 90), and its genome (B1B1B2B2B2B2) is quite complex due to polyploidy, self-incompatibility, and high heterozygosity.

In recent years, sweetpotato molecular breeding has made many advancements, including molecular markers, gene function verification, gene editing, and other aspects, although sweetpotato is still dominated by traditional crossbreeding methods. Molecular markers are widely used in genetic diversity, germplasm identification, and marker-assisted breeding. Anglin *et al*. (2021)^6^ conducted the most comprehensive genotyping of sweetpotato germplasm resources (5,979 varieties) so far, with single-sequence repeat (SSR) markers to assess genetic characteristics, diversity, and population structure. SSR markers were also used to genotype sweetpotato varieties from Africa and America^7^. To optimize the breeding process, Kumar et al. (2020)^8^ developed single nucleotide polymorphism (SNP) markers for quality assurance and control (QA/QC) based on 662 sweetpotato varieties. Through resequencing, Xiao et al. (2020)^9^ designed 40,366 pairs of indel markers covering the whole genome, among which 3,219 high-quality marker pairs were selected to construct a core marker set for sweetpotato.

More and more genes for important traits such as resistance, quality, growth, and development of sweetpotato have been discovered. Thus far, the discovered genes have been related to the salt tolerance, drought resistance, and disease resistance of sweetpotato; they include *IbP5CR, IbNFU1, IbMas, IbSIMT1, IbNHX2, IbDFR, IbMYB1, IbC3H18, IbMIPS1, IbNAC1, IbpreproHypSys, IbBBX24, IbRAP2.4, IbGATA24, IbIPUT1, IbCAR1, IbPYL8, IbbHLH66, IbbHLH118,* and others^10–20^. The improvement of sweetpotato quality has focused mainly on starch, carotenoid, and anthocyanin. At present, the main genes related to quality are *IbGBSSI, IbSBEII, IbAATP, SRF1, IbSnRK1, IbMYB1-2, IbbHLH2, IbCYP82D47, IbGGPS*, and others^21–29^. Tuberous root development is also a trait of concern in sweetpotato, and some related genes have been studied, including *IbEXP1* and *IbNAC083*^30, 31^. Genome editing is a revolutionary technology in molecular breeding. However, there are few reports on sweetpotato gene editing. Wang et al. (2019)^21^ used CRISPR-Cas9 technology to obtain a transgenic sweetpotato with high amylose and waxy starch contents, providing new raw materials for food processing and industrial applications. Yu et al. (2021)^32^ used CRISPR-Cas13 technology to introduce the *RfxCas13d* system targeting *spCSV-RNASE3* into sweetpotato, which significantly reduced the accumulation of SPCSV and SPFMV viruses in transgenic plants and improved resistance. Gene editing systems depend on a stable genetic transformation system. The embryogenic suspension line is currently the most widely used method. Recently, the genetic transformation system of sweetpotato was greatly optimized by the CDB method without tissue culture, by which the amylose content in tuberous root was changed by editing the *GBSSI, SBEI,* or *SBEII* gene^33^. Due to the complexity of genetic background, both research into and application of sweetpotato molecular breeding will face more difficulties. To solve this problem, we need a perfect sweetpotato whole genome sequence^34^.

Draft sequences of genus *Ipomoea* were first reported in *I. trifida*^35^ with Illumina short reads. Then, chromosome-scale genome sequences were reported in *I. nil*^36^, *I. trifida*^37, 38^, and *I. triloba*^37^. Of these, the 2 *I. trifida* genomes were based on primary contigs (unphased sequences) generated from heterogeneous accessions. Recently, haplotype-resolved genome sequences have been constructed in many plant species from accessions with heterogeneous genome structures^39^. Primary contig sequences, which are created from multiple heterozygous haploid genomes, are created by selecting representative sequences from multiple haploid genomes and artificially mixing them. Therefore, the obtained assembled sequences do not reflect the ‘true’ genome sequence of a sequenced individual. To compare genome structures in pan-genome analysis or to analyze the regulation of gene expression by cis- or trans- positions on the genome, it is necessary to determine the genome sequence reflecting the haploid genome, not the artificially created primary contig sequence. In sweetpotato, haplotype-resolved genome sequences were reported by Yang et al. (2017)^34^. Phasing was performed with SNP alleles identified on the scaffolds, and an ∼824 Mb assembly was created based on short reads obtained by the Illumina HiSeq 2500 and Roche GS FLX+ platforms. That was the first report of sweetpotato genome assembly; however, the total length of the assembled sequences was less than 30% of the total length of the haplotype genome.

In this study, we established a haploid-resolved and chromosome-scale *de novo* assembly of autohexaploid sweetpotato genome sequences. Before constructing the sweetpotato genome, we created chromosome-scale genome sequences in *I. trifida* using a highly homozygous accession, Mx23Hm, with PacBio RSII and Hi-C reads. The sweetpotato genome sequences were created by hybrid assembly with Illumina paired-end (PE) and mate-pair (MP) reads, 10X genomics reads, and PacBio RSII reads. Then, 90 chromosome-scale pseudomolecules were generated by aligning the scaffolds onto a sweetpotato linkage map. This is the first report of chromosome-scale *de novo* assembly of the sweetpotato genome, and the results are expected to contribute to genomic and genetic analyses of sweetpotato.

## Results and Discussion

### Genome assembly of I. trifida Mx23Hm

A total length of 64.26 Gb of PacBio subreads was obtained for an *I. trifida* inbred line, Mx23Hm, from 71 SMRT cells (Table S1). The genome size of Mx23Hm was previously estimated as 515.8 Mb^35^. Therefore, the total obtained genome sequences were estimated as 124.6x of the genome size of Mx23Hm. Illumina PE and MP reads (insert sizes = 5Kb, 15 Kb and 20 Kb) were also obtained by MiSeq and HiSeq 2000, for the genome assembly (Table S1). In addition, 64.5 Gb of PE (DRX021659), 16.3 Gb of 3 Kb inert MP (DRX21660), and 16.9 Gb of 10 Kb inert MP reads (DRX21661) obtained in the previous study^35^ were used in the assembly.

*De novo* assembly was conducted with the subreads using the SMRTMAKE assembly pipeline^40^. A total of 2,881 contigs were generated with a total length of 495.7 Mb (Table S2). The 2,881 contigs were polished with the PE reads (DRX021659) using BWA^41^ and Genome Analysis Toolkit (GATK)^42^, and scaffolded by using SSPACE 3.0^43^ with the all of the MP reads and the MiSeq PE reads. The resultant number of scaffolds was 582, with total and N50 lengths of 497.0 Mb and 10.8 Mb, respectively. Chromosome-scale scaffolding was then performed by HiRise^44^ with 471 M Hi-C reads obtained from HiSeq 2500. The numbers of breaks and joints made by HiRise were 39 and 102, respectively, and 520 sequences including 15 chromosome-scale scaffolds were created. The sequences of the 15 chromosome-scale pseudomolecules were compared with *I. nil* genome sequences (Asagao_1.2^36^), by Nucmer^45^, and were given corresponding chromosome numbers (chr01 ∼ chr15) according to those in the *I. nil* genome (Fig. S1). The chr0 sequences were then created with the remaining 505 scaffold sequences concatenated with N10000. The 15 pseudomolecules and the chr0 sequences were designated as Itr_r2.2 (Table 1, Table S2). The total length of Itr_2.2 was 502.2 Mb, including total lengths of 460.77 Mb for the 15 pseudomolecules and 41.47 Mb for the chr0 scaffold. Itr_r2.2 covered 97.4% of the Mx23Hm genome, when the genome size was considered 515.8 Mb^35^, while the cover ratio of the 15 chromosome-scale scaffolds was 89.3%.

**Table 1.** Statistics on the assembled I. trifida (Itr_r2.2) and I. batatas (IBA_r1.0) genome sequences and CDSs.

|  |  |  |  |  |  |  |  |
| --- | --- | --- | --- | --- | --- | --- | --- |
| Species and accession | *I. trifida*, Mx23Hm |  |  | *I. batatas*, Xhushu 18 |  |  |  |
| Sequence | Itr_r2.2 |  |  | IBA_r1.0 |  |  |  |
|  | Chr01~15 + Chr0 | Chr01 ~ 15 | Gene | All | Chr | Unplaced | Gene |
| Number of sequences | 16 | 15 | 34,386 | 109,986 | 90 | 109,896 | 175,633 |
| Total length of sequences (bp) | 502,237,654 | 460,770,816 | 36,202,112 | 2,907,375,442 | 2,168,449,239 | 738,926,203 | 181,148,040 |
| Average length of sequences (bp) | 31,389,853 | 30,718,054 | 1,053 | 26,434 | 24,093,880 | 6,724 | 1,031 |
| Max length of sequences (bp) | 41,466,838 | 38,817,897 | 16,284 | 40,468,515 | 40,468,515 | 6,502,119 | 20,832 |
| N50 length (bp) | 31,779,616 | 31,779,616 | 1455.00 | 23,279,505 | 26,187,252 | 17,387 | 1,380.0 |
| GC% | 37.2 | 36.5 | 46.4 | 36.1 | 35.7 | 37.2 | 46.20 |
| N% | 1.30 | 0.18 | 0.00 | 0.07 | 0.14 | 0.34 | 0.00 |
| BUSCO v5.2.2 embryophyta_odb10、 Number of BUSCOs = 1,614 |  |  |  |  |  |  |  |
| Complete | 98.5 | 98.5 | 82.6 | 99.5 | 99.4 | 56.1 | 56.1 |
| Complete single | 93.4 | 93.5 | 78.5 | 1.7 | 2.5 | 35.5 | 35.5 |
| Complete double | 5.1 | 5.0 | 4.1 | 97.8 | 96.9 | 20.6 | 20.6 |
| Fragmented | 0.8 | 0.8 | 5.1 | 0.2 | 0.2 | 6.8 | 6.8 |
| Missing | 0.7 | 0.7 | 12.3 | 0.3 | 0.4 | 37.1 | 37.1 |

To investigate the possible misassembly of Itr_r2.2, a linkage map was constructed with 190 F_1_ individuals derived from a cross between the 2 *I. trifida* accessions, 0431-1 and Mx23-4. dd-RAD-Seq sequences were obtained for the mapping population and mapped onto the Itr_r2.2 genome. A linkage map consisting of 15 linkage groups (LGs) was constructed with 8,172 variants by using LepMap3^46^ (Table S3). The linear correspondence between the physical and genetic positions of the 8,172 variants indicated few possible misassemblies at the chromosome level (Fig. S2). The total sequence length of Itr_r2.2 (502. 2Mb) was longer than those of previously published *I. trifida* genomes, V3 (492.4 Mb) and ASM470698v1 (460.9 Mb; Table S4). Meanwhile, N% of Itr_r2.2 (1.3%) was the lowest in the 3 *I. trifida* genomes. The GC% of Itr_r2.2 was 37.2%, which was higher than those of V3 (35.7%) and ASM470698v1 (35.6%).

The assembly quality of Itr_r2.2 was then investigated by mapping the sequences onto 1,614 BUSCOs (Table 1, Table S4). The results demonstrated that the ratio of complete BUSCOs was 98.5%, including 93.4% of single-copy genes and 5.1% of duplicated genes. The ratios of fragmented and missing BUSCOs were 0.8% and 0.7%, respectively. The ratios of identified complete BUSCOs on the 15 pseudomolecules were almost the same as those on Itr_r2.2, suggesting high gene coverage ratios in the 15 pseudomolecules. The BUSCO results were similar to the previously published *I. trifida* and *I. triloba* genomes, whose complete BUSCO ratios were 98.6% *(I. trifida* V3), 98.5% (*I. trifida* ASM470698v1), and 98.6% (*I. triloba* v3; Table S4).

The sequence structures of the 15 pseudomolecules in the Itr_r2.2 genome were compared with those of the *I. trifida* (*I. trifida* V3 and ASM470698v1) and *I. triloba (I. triloba* V3) genomes. High sequence homology was observed across the entire regions between Itr_r2.2 and the 3 genomes (Fig. S3). However, several differences were also observed. For example, partial sequence inversions were observed on chr02, chr03, chr04, chr07, and chr08 between Itr_r2.2 and *I. trifida* V3. The obvious structural differences were smaller in the genome comparison between Itr_r2.2 and ASM470698v1 than between Itr_r2.2 and *I. trifida* V3. However, the genome-wide sequence similarity between Itr_r2.2 and ASM470698v1 seems lower than that between Itr_r2.2 and *I. trifida* V3. These results suggested that the 3 *I. trifida* genomes had different genome structures. Interestingly, the genome structures between *I. triloba* v3 and Itr_r2.2 were more similar than those between *I. trifida* v3 and Itr_r2.2 in many chromosomes. However, in chr01, for example, sequence similarity was quite low and was observed only at both ends of the chromosome. It was difficult to determine whether the differences in sequence structure were real or only the result of poor assembly quality. Meanwhile, Itr_2.2 was an S_11_ descendant of a heterozygous accession, Mx23-4^35^, and therefore may more accurately reflect the genomic structure of the sequenced material than the other 2 genomes, which were constructed as unphased genomes. Moreover, among the genomes Itr_r2.2 was the longest in total length and the lowest in N%. We therefore concluded that Itr_r2.2 was suitable as a reference for sweetpotato genome assembly.

### Sweetpotato genome assembly

Whole genome shotgun sequences of a sweetpotato variety, Xushu 18, were obtained from the Illumina PE, MP, and 10X Chromium Genomics libraries (Table S1). A Smudgeplot (K-mer = 21)^47^ based on 252.2 Gb of 250 nt PE reads (DRX405125) indicated that the most likely genome structure of Xushu 18 was an auto-hexaploid, AAAAAB (Fig. S4). The haploid genome size of Xushu 18 was estimated as 294.6 Mb by using GenomeScope2.0 (k-mers = 19, hexaploid mode; Fig. S5)^47^. The ratio of heterozygous sequences on the genome was estimated as 7.31%. The ratio of unique sequences was estimated as 45.6%. Because GenomeScope2.0 seems to underestimate genome size in polyploid species, the genome size of Xushu 18 was investigated also on the basis of the distribution of distinct k-mers (K = 17) identified by Jellyfish^48^ with a total length of 215.7 Gb of 151 nt PE reads (random sampling of DRX409453). One large and 2 small peaks were observed in the distribution plot, and the genome size of Xushu 18 was estimated as 2,603 Mb based on the large peak at multiplicities of 74 (Fig. S6).

The results of genome size estimation varied in previous studies. For example, Ozias-Akins and Jarret (1994)^49^ reported that the 2C content of the sweetpotato nucleus was 4.8–5.3 pg/2C, while Srisuwan et al. (2019)^50^ reported 3.1-3.3 pg/2C. Given that the haploid genome size of the diploid *I. trifida* haploid is around 500 Mb, it is reasonable to assume that the genome size of sweetpotato is around 3 Gb/2C. We considered it possible that Jellyfish (2.6 Gb) led to an underestimation due to the influences of homologous sequences across homoeologous chromosomes.

In addition to the short reads, a total length of 181.5 Gb of PacBio CCS reads was obtained from 206 SMRT cells. Denovo MAGIC 3 (NRGene, Ness Ziona, Israel) was used for assembly, and 110,708 scaffolds and 1,567,871 unplaced contig sequences were created (Table S5). The total length of scaffolds was 2,907.4 Mb, while that of unplaced contigs was 575.7 Mb. As the sequence lengths of the unplaced contigs were very short, with an N50 length of 451 bp, only the scaffold sequence was subjected to further analysis.

To create chromosome-scale scaffolds, an S_1_ linkage map was constructed using the variants identified on the *I. trifida* genome. The dd-RAD-Seq sequences of 437 S_1_ individuals were mapped onto 520 scaffolds comprising the Mx23Hm Hi-C scaffolds (Table S2). To construct a linkage map, a total of 28,516 variants that showed simplex segregation, i.e., that fit the expected ratio of 1:2:1 via Chi-square tests (P[≥[01), in the population were used. The linkage map was constructed by using Onemap^51^, and a total of 27,858 variants were mapped on 96 linkage groups (LGs) (Tables S6 and S7). The 50 bp sequences on both sides of the mapped variants on the Mx23Hm Hi-C scaffolds were then cut out and corresponding positions on the cut-out sequences on the 110,708 Xushu 18 scaffolds were identified by a BLAST analysis. This identified a total of 22,537 variants corresponding to positions on 959 Xushu 18 scaffolds. Of the 959 scaffolds, 534 were aligned with a unique corresponding LG, while the remaining 425 corresponded to positions across multiple LGs. In the latter cases, the scaffolds were cut and aligned by ALLMAPS^52^.

The aligned sequences on the 96 LGs of the S_1_ linkage maps were compared with Itr_r2.2 and given chromosome numbers. Homologous chromosomes were identified by the lowercase letters a-f, with the sequence most similar to that of *I. trifida* designated as “a” and the sequence most different as “f” (Fig. S7). Two pairs of sequences, aligned on lb11-6 and lb11-7 and on lb14-6 and lb14-7, were combined and given the chromosome IDs chr01f and chr03f, respectively (Table S6, Table S7). Because of their short lengths, the sequences aligned on lb02-7, lb02-8, lb07-7, and lb07-8 were classified as unplaced scaffolds. As a result, a total of 90 chromosome-level pseudomolecules and 109, 896 unplaced scaffolds were created and were designated as IBA_r1.0 (Table 1, Table S5). The genome structure of sweetpotato was compared with that of *I. trifida* (Itr_r2.2), and entire syntenic relationships were observed across the 90 pseudomolecules (Fig. 1).

**Fig. 1.**
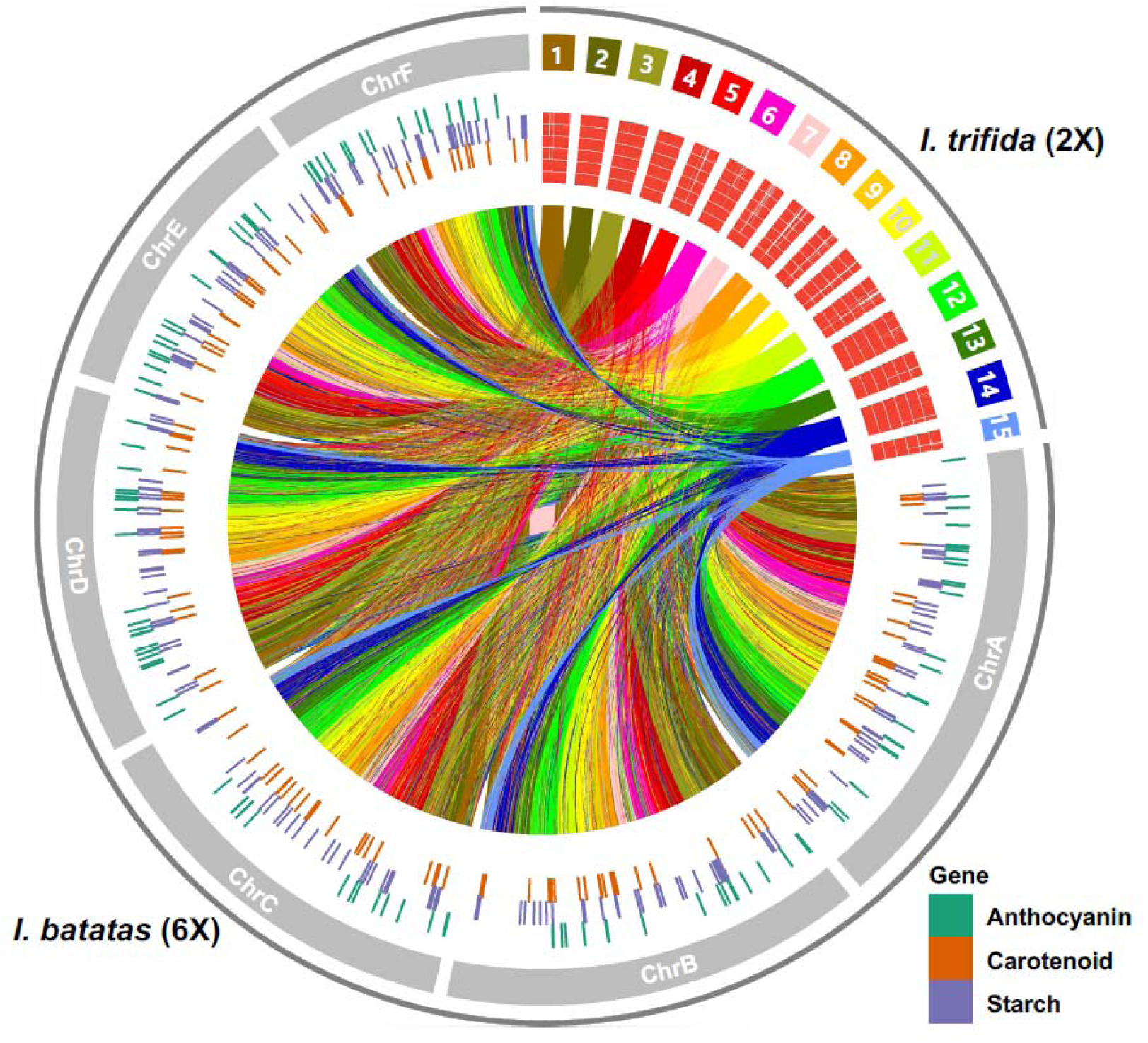
Genome sequence comparison between *I. trifida* Itr_r2.2 and sweetpotato (*I. batatas*) IBA_r1.0. Gene locations relating to anthocyanin, carotenoid, and starch synthesis are represented by green, blue, and red bars, respectively. The Red blocks under the 15 chromosomes of I. trifida shows gene regions paired with *I. batatas.* If there are 6 tiles (bars), it means that the same genes are distributed on all 6 homoeologous chromosomes.

The total length of IBA_r1.0 was 2,907.4 Mb, consisting of 2,168.4 Mb pseudomolecules and 738.9 Mb unplaced scaffolds (Table 1). The 90 pseudomolecules and the 109,896 unplaced scaffolds occupied 74.6% and 25.4% of IBA_r1.0, respectively. The lengths of the 90 pseudomolecules ranged from 40.5 Mb (chr12a) to 9.2 Mb (chr15f; Table S8) with a mean length of 24.1 Mb. The ratio of complete BUSCOs assembly on IBA_r1.0 was 99.5%, including 1.7% of single-copy genes and 97.8% of duplicated genes. The ratio of complete BUSCOs identified on the 90 pseudomolecules was almost the same as that on all Itr_r2.2, 99.4%, including 2.5% of single and 96.9% of duplicated genes. The high ratios of complete BUSCOs suggested the assembled sequences covered most of the gene sequences on the Xushu 18 genome.

The assembly quality in IBA_r1.0 was investigated by using Merqury^53^ (Fig. S8). The completeness and QV were estimated as 96.91% and 25.15, respectively. The error rate was 0.00305. K-mer multiplicity plots were clearly observed for all copy number plots of x1, x2, x3, x4, and x >4 copies; as Xushu 18 was a hexaploid, each copy number plot reflected the degree of homology between the homologous chromosomes. Two peaks were observed in the x2 and x3 plots, with the left-hand peak overlapping the x1 or x2 peak, respectively. This suggested a possibility that partial duplicate or triplicate sequences in the Xushu 18 genome artificially collapsed during the assembly process. The x4 and >4 copy plots showed no clear peak separation, but the plots overlapped with the x1 ∼ x4 peaks, suggesting the artificial collapse of the assembled sequences was also happened. The ratios of genome sequence reads (DRX405125) classified as x1, x2, x3, x4, and x >4 copy were calculated based on the numbers obtained by multiplication of k-mer multiplicities and counts (Fig. S9). The ratios of sequences in x1, x2, x3, and x4; that is, single, double, triple, and tetra homoeologous chromosome sequences, ranged from 6.9% to 8.0%. Meanwhile, 70.8% of reads were classified as x >4, which was considered to be conserved in 5 or 6 homoeologous chromosomes.

### Chloroplast genome and mitochondrial genome assembly

Chloroplast genomes of *I. trifida* (Mx23Hm) and *I. batatas* (Xushu 18) were assembled with about 1.5 Gb MiSeq and 453 Mb HiSeq reads, respectively, based on the reference, the initial chloroplast genome of *I. batatas* (KP212149). The assembled chloroplast genome sizes were 161,693 bp (the average depth is 196X) and 161,429 bp (80x), respectively, with no gaps. The GC content of each chloroplast genome was the same, 37.53% (Table S9). The chloroplast genomes contained 112 genes, and the genes were classified into 78 protein coding genes, 30 tRNA genes, and 4 rRNA genes (Table S10). The gene maps of the chloroplast genomes were constructed by the OGDRAW program (Fig. S10). Using MEGA11, a phylogenetic tree for 17 species was constructed with 68 common protein-coding sequences in the chloroplast genomes (Fig. S11)^54^. The tree shows that Mx23Hm and Xushu 18 have the closest relationships with *I. tabascana*. Xiao et al. (2021)^55^ analyzed the chloroplast genomes of 107 sweetpotato cultivars and reported that the average GC content was 37.54% and that sequence homology between them was higher than 98%. The GC contents in the chloroplast genomes of Mx23Hm and Xushu 18 were both 37.53%, almost the same as in the previous study.

The mitochondrial genomes of *I. trifida* (Mx23Hm) and *I. batatas* (Xushu 18) were assembled with 10.1 Gb and 11.4 Gb MiSeq reads, respectively, based on the *I. nil* mitochondrial genome (AP017303). The complete mitochondrial genome lengths of Mx23Hm and Xushu 18 were 264,698 bp and 269,586 bp, respectively. The mitochondrial genomes of Mx23Hm and Xushu 18 had 55 and 57 predicted genes, respectively. Both genomes had 34 protein-coding genes and 3 rRNA genes. However, the numbers of tRNA genes differed: 18 in Mx23Hm and 20 in Xushu 18. Resulting in a difference in the total number of genes. The gene map of the mitochondrial genomes drawn by the OGDRAW program shows the different distributions of the genes between the 2 genomes (Fig. S12). Using MEGA11, phylogenetic analysis based on the mitochondrial genome was performed with 9 species using 10 representative protein-coding sequences (Fig. S13). The phylogenetic analysis shows that Mx23Hm and Xushu 18 had the closest relationship. The high-quality organelle genomes of Mx23Hm and Xushu 18 can be used widely to increase the accuracy of genetic structures and for evolutionary studies.

### Repetitive sequences

To analyze how repeats affect sweetpotato genome sizes, repeats present in the genomes of 502 Mb of *I. trifida* Mx23Hm (ITR_r2.2) and 2.9 Gb of *I. batatas* Xushu 18 (IBA_r1.0) were predicted. RepeatModeler^56^ was used to create a custom database of each genome assembly independently, and Repbase^57^ was used to annotate these custom libraries. Genome assemblies were masked using RepeatMasker^58^ with the custom libraries. As a result, 278 Mb (55.44%) in *I. trifida* and 1.5 Gb (52%) in *I. batatas* were masked as repeat regions (Table S11). Concretely, annotated repetitive sequences have higher ratios of RNA elements than DNA elements. The LTR regions of *I. trifida* and *I. batatas* were 17.66% and 15.97%, respectively. Repetitive elements are inserted in every compartment of the genomes and participate in the expansion of the intronic regions in *I. batatas* and *I. trifida*. In the case of small RNA, the repeat proportions were 1.76% in *I. trifida* and 0.39% in *I. batatas*, showing a significant difference between the 2 cultivars.

In the haplotype-resolved *I. batatas* Taizhong 6 genome, the total repeat was 45.6% and the LTR element showed a relatively low ratio of 10.9%^34^. On the other hand, in the case of *I. nil,* the total repeat was 63.29% and the LTR element was 21.68%, showing a higher tendency than other species^36^. *I. trifida* and *I. triloba* showed 50.2% and 52.8% of repeat sequences^37^. The repeat sequence ratios in the two *Ipomoea* genomes assembled in this study showed a similar trend to those in the previous studies. Thus, the difference in genome size between *I. trifida* Mx23Hm and *I. batatas* Xushu 18 can be attributed mostly to polyploidy.

### Gene prediction and functional annotation

The schematic workflow for protein coding gene prediction is shown in Fig. S14. For gene prediction in Itr_2.2, a consensus gene model was created based on *ab initio* gene models created by Augustus, evidence-based gene models based on transcript sequences derived from 5 tissues in Mx23Hm and 3 tissues in Xushu18, and 5 coding gene models derived from *Solanum tuberosum*^59^, *S. lycopersicum*^60^, *Manihot esculenta*^61^, *Oryza sativa*^62^, and *Arabidopsis thaliana*^63^ (Table S12). For gene prediction in IBA_r1.0, transcript sequencing was conducted for 6 tissues in Xhushu 18 with Illumina PE transcript sequences (Table S1). The previously obtained transcriptome sequences were also used for gene prediction (Table S13). Evidence-based gene prediction was also performed with gene models derived from *I. trifida*^37^, *I. triloba*^37^, *I. tabascana* (unpublished, derived from Sweetpotato Research Institute, CAAS), and *I. nil*^36^ as well as *S. tuberosum*, *S. lycopersicum*, *M. esculenta*, *O. sativa*, and *A. thaliana* (Table S12). Sequences coding transcription factors in plants were also used by reviewing PlantTFDB. The predicted gene sequences were merged with those created by Augustus.

Specifically, for Itr_r2.2 gene prediction, 34,386 consensus gene sequences were created with 48,701 gene sequences from Augustus, 137,471 gene sequences predicted from transcript sequences, and 26,080 ∼ 63,0385 gene sequences from protein coding gene models in 5 plant genomes (Table 1, Table S14). Meanwhile, for IBA_r1.0 gene prediction, 175,633 consensus gene sequences were created with 255,611 gene sequences from Augustus, 1,026,264 gene sequences predicted from transcript sequences, and 66,945 ∼ 525,770 gene sequences from protein-coding gene models from 10 plant genomes and PlantTF (Table 1, Table S15).

The total length of the genes in Itr_r2.2 was 96.3 Mb, whereas that in IBA_r1.0 was Mb (Table 1, Table S16). The gene coverages on the genomes were similar in both genomes: 19.2% in Itr_r2.2 and 19.0% in IBA_r1.0. The quality of gene prediction was assessed by BUSCO analysis. In Itr_r2.2, the ratio of complete BUSCOs was 82.6%, including 78.5% of single-copy genes and 4.1% of duplicated genes, whereas in IBA_r1.0 the ratio was 97.2%, including 9.5% of single-copy genes and 87.7% of duplicated genes. It was considered that the higher number of predicted genes in IBA_r1.0 was caused by duplicate gene sequences located across homoeologous chromosomes. The average number of exons per gene was 4.4 to 4.3, and the average length of one exon ranged from 236.4 bp to 242.6 bp (Table S17, Fig. S15). While the average length of genes was slightly longer in IBA_r1.0 (3,136.5 bp) than in Itr_r2.2 (2799.9 bp), the number of exons per gene was similar between the 2 genomes, suggesting similar gene structures between them. Interestingly, the GC content of Ipomoea species ranged from 35.6% to 36.7%, higher than the 30.59% of *M. esculenta*, 28.43% of *S. tuberosum*, and 30.49% of *S. lycopersicum*.

The functional annotation of the predicted genes was conducted with BLAST search and gene ontology (GO) analyses using Blast2Go software^64^ and Trinotate^65^ against 7 public databases: NCBI nr^66^, UniProt^67^, GO^68^, KEGG^69^, Pfam^70^, SignalP^71^, and TmHMM^72^. In the case of Itr_r2.2, 4,093 (11.9%) of genes had no similar genes in the databases (Table S18). However, the other genes were functionally annotated, as follows: nr BLAST 29,767 (86.57%), UniProt 23,109 (67.20%), GO 23,783 (69.16%), KEGG 19,919 (57.93%), Pfam 21,387 (68.20%), SignalP 2,964 (8.62%), and TmHMM 6,497 (18.89%; Table S18). Meanwhile, 170,317 (96.97% of the total) genes on IBA_r1.0 were annotated by BLAST analysis against NCBI nr (85.42%) and UniProt (80.90%) databases. The functions of most of the annotated genes were predicted based on *I. nil* gene annotation. The numbers of annotated genes by GO, KEGG and Pfam were 105,620 (60.14%), 57,735 (32.87%), and 91,519 (52.11%), respectively. SignaIP and TmHMM gave functional annotation to 16.925 (9.64%) and 28,348 (16.14%) genes.

Transcription factors regulate gene expression by binding to sequence-specific DNA-binding proteins, and the expression of transcription factor genes affects their phenotypic traits. Transcription factor genes of *I. trifida* Itr_r2.2 and *I. batatas* IBA_r1.0 were predicted by InterProScan^73^ with the TF family assignment rules of PlanTFDB v5.0 (http://planttfdb.gao-lab.org/). In Itr_r2.2 and IBA_r1.0, 1,327 and 7,744 transcription factor genes were predicted, respectively. The compositions of the transcription factor genes among the 8 plant species were compared, revealing that the MYB superfamily showed the highest number of genes among all of the compared species except *I. nil*, in which the FAR1 transcription factor gene was the most common (Table S19).

### Comparison with other species at gene level

Comparative genomics is the study of the identification of functional associations among selected species. While comparing the genomes of selected species, the sequence similarities among them could decide the relationship distances in evolutionary studies. To study the evolutionary characteristics of sweetpotato in *Ipomoea*, we conducted a comparative genome analysis using a total of 9 genome sequences, including 5 *Ipomoea* species, 2 *Solanacea* species, and 2 *Rosids* species. First, the ortholog gene families and species-specific gene families were obtained through ortholog gene analysis (Fig. S16, Table S20). In all species, more than 80% of the genes were included in ortholog groups, and 0.51 to 8.85% of the genes were species-specific. In particular, diploid species had an average of 1.34 to 1.90 genes per ortholog group, whereas *I. tabascana*, a 4X species, contained an average of 2.24 genes, and *I. batatas*, a 6X species, contained an average of 3.48 genes per ortholog group.

### Starch, anthocyanin, and carotenoid pathway genes

Starch, anthocyanin, and carotenoid pathway gene analysis studies provide available information for research into the isolation of useful trait genes and the breeding of sweetpotato. We used the CAZY^74^ and KEGG^69^ pathway databases to identify genes related to starch, anthocyanin, and carotenoid biosynthesis. After downloading related genes from the database, we predicted the genes by mapping them to the *I. trifida* and *I. batatas* genome sequences. To find genes related to the anthocyanin and carotenoid pathways, additional analysis was performed using anthocyanin and carotenoid gene information from the NCBI database. As a result of the prediction, we identified 216 starch pathway-related genes in 52 family groups (Table S21), 175 anthocyanin genes in 37 family groups (Table S22), and 162 carotenoid genes in 35 family groups (Table S23) in *I. batatas*. In the case of *I. trifida*, we identified 45 genes in 45 family groups for the starch pathway, 41 genes in 35 family groups for the anthocyanin pathway, and 34 genes in 33 family groups for the carotenoid pathway.

The distribution of genes in the *I. batatas* genome shows the relationship between the genes’ physical locations on the genome and the expression of traits (Figure S17). Some of the starch, anthocyanin, and carotenoid genes were located close to each other. Gemenet et al. (2020)^75^ reported that the contents of beta-carotene and starch are negatively correlated in sweetpotato due to the physical linkage of the phytoene synthase gene and the sucrose synthase gene. In this study, we found that the phytoene synthase gene (Iba_chr02aCG5430, Iba_chr02cCG4140, Iba_chr02fCG4060,) and the sucrose synthase gene (Iba_chr02aCG5440, Iba_chr02cCG4130, Iba_chr02fCG4050) were adjacent to each other in the *I. batatas* Xushu 18 genome (Tables S21, S23).

For the expression analysis of starch, anthocyanin, and carotenoid genes in *I. batatas* tissues, RNA was isolated from the leaves at 42 days after transplantation (DAT), stems at 42 DAT, and roots at 90 DAT. RNA sequencing (RNA-seq) was performed using a strand-specific library. The trimmed RNA-seq was attached to the Xushu 18 genome, and the fragments per kilobase of transcript per million mapped reads (FPKM) values were calculated and used for expression profiling of genes. A heatmap was generated with the average FPKM values in 2 replicates using the R-package pheatmap (ver. 1.0.12).

The expression heatmap showed that starch pathway genes of sucrose synthase 1a, sucrose synthase 1b, ADP-glucose pyrophosphorylase L1, ADP-glucose pyrophosphorylase L2 ADP-glucose pyrophosphorylase S1, granule-bound starch synthase 1, starch synthase 2a, starch branching enzyme 2, starch phosphorylase, UDP-glucose pyrophosphorylase, β-amylase 1, and β-amylase 3 genes were highly expressed in root; sucrose synthase 1, ADP-glucose pyrophosphorylase L3, and starch debranching enzyme 1 genes were highly expressed in stem; and ADP-glucose pyrophosphorylase L4 was highly expressed in leaf (Fig. S18, Table S24A).

High expression levels were observed in leaf for the anthocyanin pathway genes (Fig. S19, Table S24B) including phenylalanine ammonia lyase 2, phenylalanine ammonia lyase 5, cinnamic acid 4-hydroxylase, 4-coumarate CoA ligase, chalcone synthase, flavanone 3- hydroxylase, flavonoid 3’-hydroxylase, anthocyanidin 5-O-glucosyltransferase 2, and carotenoid pathway genes (Fig. S20, Table S24C), including geranylgeranyl pyrophosphate synthase S1, phytoene synthase, phytoene desaturase, lycopene beta cyclase, zeaxanthin epoxidase, violaxanthin de-epoxidase, beta carotene isomerase, and carotenoid cleavage dioxygenase genes. The results showed that starch biosynthesis pathway genes were expressed in root and that genes of the anthocyanin and carotenoid biosynthesis pathways were expressed in leaf. Many studies have been conducted on the starch^76^, anthocyanin^77^, and carotenoid biosynthesis pathways^78, 79^ in plants. However, few such studies have been conducted on sweetpotato. Thus, the structural and expression analysis results in this study will contribute greatly to understanding how those genes work and function in the starch, anthocyanin, and carotenoid biosynthesis pathways in sweetpotato.

### T-DNA regions

T-DNAs in the sweetpotato genome, such as the T-DNA of *Agrobacterium rhizogenes* in the domesticated sweetpotato, result from a natural horizontal gene transfer (HGT) by the *Agrobacterium* genus^80^. In the sweetpotato genome, *Ib*T-DNA1 (GenBank accession: KM052616; 10829 bp) originating from an unknown *Agrobacterium* species and *Ib*T-DNA2 (GenBank accession: KM052617; 12,075 bp) originating from *A. rhizogenes* were found based on sequence similarity. Concretely, *Ib*T-DNA1 on chromosome 12 had 4 copies: 2 copies in chr12a and 1 copy each in chr12b and chr12c (Table S25A). In contrast, *Ib*T-DNA2 had 4 copies at the 50.76 kb region of chromosome 7d (Iba_chr07d: 99449-15210, 50,762 bp) (Table S25B). These results suggest that in the case of *Ib*T-DNA1, horizontal gene transfer of T-DNA is followed by whole genome triplication. This, in turn, is followed by a tandem duplication event for the T-DNA on chromosome 12a. On the other hand, in the case of *Ib*T-DNA2, the result suggests that horizontal gene transfer to the chromosome 7d 50.76 kb region event occurred after whole genome triplication, and that the tandem duplication occurred later.

The *Ib*T-DNA1 region contains 2 tryptophan-2-monooxygenases (iaaM), 3 indole-3- acetamide hydrolases (iaaH), 3 C-proteins, and 6 agrocinopine synthase (Acs) genes (Table S73). In the IbT-DNA2 region, 2 ORF17n, 5 rooting locus RoolB/RolC, 3 ORF13, and 4 ORF18 gene copies were identified (Table S26). We did not find the *Ib*T-DNA sequences in diploid sweetpotato *I. trifida* Mx23Hm or in tetraploid sweetpotato *I. tabascana* Ganshu (unpublished). The results suggested that the ancestor of the hexaploid sweetpotato underwent a different T-DNA insertion event than the other *Ipomoea* species. Kyndt et al. (2015)^80^ analyzed T-DNA region sequences with sweetpotato var. Huachano. The T-DNA copy number of Huachano was tested through Southern blot analysis, revealing that *Ib*T-DNA1 showed 4 to 6 bands and that *Ib*T-DNA2 showed 4 bands. These results were consistent with the 4 copies of *Ib*T-DNA1 and 4 copies of *Ib*T-DNA2 present on the genome of Xushu 18 that we assembled.

## Methods

### Plant materials

Genome assembly and transcriptome analysis in *I. trifida* were performed using accession Mx23Hm, developed at the Kyushu Okinawa Agricultural Research Center, National Agriculture and Food Research Organization (KOARC, NARO), Japan. Mx23Hm is a single descendant selfed line (S_11_) derived from the self-compatible experimental line Mx23-4, which was introduced from Mexico to Japan in 1961. A linkage map was constructed with 190 F_1_ individuals derived from a cross between 2 *I. trifida* accessions: 0431-1 and Mx23-4. For genome assembly and transcriptome analysis in *I. batatas*, the Chinese variety Xushu18, bred in Xuzhou Institute of Agricultural Sciences in Jiangsu Xuhuai District in 1977, was used. The linkage map was constructed with 437 S_1_ individuals derived from self-crosses of Xushu 18.

To obtain PacBio CCS reads, genomic DNA was extracted by a modified CTAB method. For short reads sequencing using Illumina platforms, DNAs were extracted by using the Genomic DNA Extraction Column (Favorgen Biotech Corp., Ping-Tung, Taiwan). Total RNA of Mx23Hm was extracted from young and old leaves, petioles, roots, and stems from materials grown in a greenhouse at KOARC, NARO, according to the protocol from Takahata *et al*. (2010)^81^.

For the transcriptome analysis in *I. batatas*, the cuttings of Xushu18 plants at the completion of the third leaf expansion were grown under a controlled environment at 25 ± 3[in a 16/8 h light/dark cycle. The leaf and stem samples were collected at 6 weeks after transplantation. At 3 months of cultivation, storage roots (> 15 mm in diameter) were harvested. Samples were harvested with 2 biological repeats. The sampled tissues were immediately frozen in liquid nitrogen and stored at -70[until further use.

Total RNA was extracted using the RNeasy Plant Mini Kit (Qiagen, Hilden, Germany) with 100 mg of plant samples. The quality of the RNA samples was verified with an Agilent 2100 Bioanalyzer (Agilent Technologies, Santa Clara, CA, USA). All samples had 28S:18S ratios in the range of 1.8–2.0 with intact 28S, 18S, and 5S RNA bands, high RNA purity, and mean RNA integrity numbers (RINs) of 7.7 ∼ 9.5, which satisfied the requirements for library construction and sequencing.

### Genome sequencing and assembly of I. trifida Mx23Hm

To prepare the library 5 μg of gDNA was used as the input material. A SMRTbell library was constructed with SMRTbell™ Template Prep Kit 1.0 according to the manufacturer’s instructions (Pacific Biosciences, Menlo Park, CA, USA). Fragments smaller than 12 Kb of the SMRTbell template were removed using the Blue Pippin Size selection system to construct a large-insert library. The constructed library was validated by the Agilent 2100 Bioanalyzer. After a sequencing primer was annealed to the SMRTbell template, DNA polymerase was bound to the complex using DNA/Polymerase Binding Kit P6. This SMRTbell adaptor complex was then loaded onto the SMRT cells. The SMRTbell library was sequenced with 71 SMRT cells (Pacific Biosciences) using C4 chemistry (DNA Sequencing Reagent 4.0), and a 1[240-minute movie was captured for each SMRT cell using the PacBio RS II (Pacific Biosciences) sequencing platform.

WGS reads were obtained from the Illumina PE libraries, whose expected inert size was 500 bp, and MP libraries, whose expected inset sizes were 5, 15, and 20 K (Table S1). The PE library was sequenced by using MiSeq (Illumina, San Diego, CA, USA) with 301 nt, while the MP libraries were sequenced by HiSeq 2000 with 101 nt. In addition, PE reads and MP reads (insert sizes 3 Kb and 10 Kb) obtained in the previous study^35^ were used in this study (DRX021659 - DRX021661).

*De novo* assembly was conducted using the SMRTMAKE assembly pipeline^40^. As the estimated genome size was 515.8Mbp and the average coverage of filtered subreads was about 112.63X for 58,095,589,571 bp (Minimum Subread Length 50 bp, Minimum Polymerase Read Quality 0.75, and Minimum Polymerase Read Length 50 bp). The first steps were to filter the reads (Options --filter=’MinReadScore=0.80, MinSRL=500, MinRL=100’) and to perform an error correction (CUTOFF option setting with GENOME_SIZE 512Mb * 30). In the next step, the Celera Assembler was used to generate a draft assembly using the error-corrected reads. Before the final assembly using the quiver algorithm, the draft assembly was polished using the quiver algorithm with the PE reads to correct errors using BWA^41^ and GATK^42^. Scaffolding was performed with the PE and MP reads to upgrade the genome assembly using SSPACE 3.1^43^ with categories B, C, and D of mate pair sequences, which were categorized by NextClip ver1.3^82^. Chromosome-scale scaffolding was then performed by HiRise^44^ with Hi-C reads obtained from HiSeq 2500 (Illumina).

To investigate possible misassembly, an F_1_ linkage map was constructed with dd-RAD- Seq sequences obtained for the 190 F1 individuals derived from a cross between 0431-1 and Mx23-4. A dd-RAD-Seq library was constructed according to Shirasawa et al. (2016)^83^, and sequencing was conducted by using HiSeq 2000 (Illumina). A variant call was performed by bcftools 0.1.19 in Samtools^84^. Segregation linkage maps were constructed by using Lep-MAP3^46^. Assembly quality was also assessed by BUSCOs v5.0^85^.

### Genome sequencing and assembly of I. batatas Xushu 18

Whole genome shotgun sequences of sweetpotato variety Xushu 18 were obtained from Illumina PE, MP, and 10X Chromium Genomics libraries by using Illumina HiSeq2500 (Table S1). PacBio CCS reads were also obtained using the same method as for *I. trifida*. Genome structure and size were estimated byScope2.0 and Smudgeplot^47^ as well as Jellyfish^48^.

*De novo* whole genome assembly was performed by using Denovo MAGIC 3 (NRGene, Ness Ziona, Israel) with PE, MP, 10X Genomics, and PacBio CCS reads. To create chromosome-level scaffolds, a Xushu 18 S_1_ linkage map was constructed with dd-RAD-Seq sequences. Library construction, sequencing, and variant calls were performed using the method described by Shirasawa et al. (2017)^86^. A linkage map was constructed by using Onemap^87^, and the assembled scaffolds were aligned on the linkage map by using ALLMAPS^52^. Assembly quality was assessed by BUSCOs v5.0^85^ and Merqury^53^. BLAST v2.2.28+ and MCScanX were performed to determine the synteny between *I. balatas* and *<u>I.trifida</u>*. All-to-all BLAST was performed and significant hits were then filtered with a cutoff of e-value 1e^-50^.

### Chloroplast and mitochondrial genome assembly

The chloroplast genome was assembled using the method of Kim et al. (2015)^88^. In brief, high-quality raw reads with Phred scores above 20 were obtained from raw sequencing reads using the CLC quality trim tool in CLC Assembly Cell package ver. 4.2.1 (Qiagen). *De novo* assembly was then implemented by the CLC Genome Assembler in CLC Assembly Cell package ver. 4.2.1 (Qiagen) with the following parameters: read distance 150 - 500 bp, similarity 0.8, length fraction 0.5. Using Nucmer, the putative chloroplast contigs were selected from a comparison with the *I. batatas* (KP212149) chloroplast genome sequence as a reference^45^. Selected chloroplast contigs were merged into a single contig through junction confirmation between contigs and inner gap filling by read mapping using the CLC read mapper with the parameters of similarity 0.8 and length fraction 0.5.

The mitochondrial genome was assembled using a method similar to that used for the chloroplast genome except that the *I. nil* (AP017303) mitochondrial genome sequence was used as a reference to select mitochondrial contigs. In addition, manual curation through BLAST searches against the NCBI nr database was performed to prevent the misassembly caused by chloroplast-derived sequences with high coverage.

Genes on the chloroplast and mitochondrial genomes were primarily annotated according to the method reported by Alverson (2010)^89^ using DOGMA (http://dogma.ccbb.utexas.edu/) and Mitofy (http://dogma.ccbb.utexas.edu/mitofy/). Ambiguous gene positions were manually verified through NCBI BLASTN searches and adjusted by manual curation using the Artemis annotation tool^90^. A circular map was drawn using OGDRAW (http://ogdraw.mpimp-golm.mpg.de).

Phylogenetic analysis was performed by the following method. Protein-coding sequences conserved among the chloroplast genomes or mitochondrial genomes of related species were identified and concatenated for I. trifida and I. batatas. The concatenated sequences were multiple-aligned using MAFFT ver. 7 (http://mafft.cbrc.jp/alignment/server/index.html)^91^ with default parameters and were used as input data for phylogenetic analysis. Phylogenetic analysis was performed using MEGA 11^54^ with the maximum-likelihood method and 1000 bootstrap replicates.

### Repetitive sequences

Repetitive sequences on the assembled *I. trifida* and *I. batatas* genome sequences were detected by RepeatModeler (http://www.repeatmasker.org/RepeatModeler.html) and RepeatScout^56^. Those predicted repeats were masked using RepeatMasker^58^ before annotation. Here RepeatModeler used reference genomes in the Repbase database (http://www.girinst.org/repbase/); those genomes were closely related to the genome as a training set, to predict the repeat regions from the genome scaffolds with the default parameters.

### Gene prediction and annotation

Protein coding genes were predicted by *ab initio* prediction, evidence-based prediction using transcript sequences, and homology-based methods. Finally, all gene models were integrated to create the consensus gene model as a final gene set for the *I. trifida* and *I. batatas* genomes. A homology search was performed for the protein coding gene sequences predicted on the genomes listed in Table 12 against *I. trifida* and *I. batatas* genomes using TBASTN (E-value ≤ 1E-4). Then the homologous genome sequences were aligned to matched proteins using Exonerate^92^ to predict the accurate spliced alignments.

Transcript sequences were obtained from RNAs described in *Plant Materials*. To construct a strand-specific library, the TruSeq Stranded Total RNA Sample Preparation Kit and ScriptSeq v2 RNA-seq library preparation kit (Illumina) were used. Constructed libraries were sequenced using the Illumina HiSeq 2500 platform. In addition to the transcript sequences obtained in this study, previously published sequences were used for the gene prediction of the Xushu 18 genome (Table S13). The transcript sequences were aligned on the assembled genomes using TopHat^93^ and PASA (http://pasapipeline.github.io/), with default parameters to predict the transcript structures and splice junctions.

*Ab initio* prediction was conducted with Augustus^94^ by using the complete transcriptome as a training matrix for HMM^72^, which was produced by PASA by using transcript sequences. Finally, all of the gene models were used to build the consensus gene model. These genes were subjected to functional annotations such as NCBI-nr^66^, UniProt^67^, GO^68^, and KEGG^69^ by using Blast2GO^64^ with the default parameters. Based on the Pfam^70^ obtained from InterProScan^73^, transcription factor genes were predicted according to the TF family assignment rules of PlanTFDB v5.0 (http://planttfdb.gao-lab.org/).

### Comparative genome analysis

Ortholog gene analysis was conducted with 9 crop plants. The family genes were assigned by OrthoMCL^95^ based on sequence similarity. The database was constructed with protein coding genes in the genomes of *I. batatas*, *I. trifida*, *I. tabascana*, *I. nil*, *I. triloba*, *S. lycopersicum*, *S. tuberosum*, *M. esculenta*, and *A. thaliana* (Table S12), and was used in a self-query using BLASTp (e-value ≤ 1e-5). The BLASTp search output was subjected to analysis using OrthoMCL with the default parameters in order to construct orthologous gene families.

### Expression profiling of starch, anthocyanin, and carotenoid pathway genes

Low-quality and duplicated reads and adapter sequences were removed from Illumina transcript sequences using Trimmomatic ver. 0.39^96^ with the default parameters. The sequences contaminated by viruses, bacteria, or humans were removed using BBDuk program ver. 38.87 (https://sourceforge.net/projects/bbmap/) with the k-mer 31 parameter. The clean, high-quality transcript sequences were mapped on the assembled sweetpotato genome (IBA_r1.0) using HISAT2 ver. 2.2.1^97^. The mapped transcript sequences on protein coding gene sequences were counted using HTSeq-count ver. 2.2.1^98^. FPKM values were calculated using the number of transcript sequence reads mapped on the protein coding gene sequences and were used for the expression profiling of genes. To visualize gene expression peaks among the different samples, a heatmap was generated with the average FPKM values in 2 replicates of genes using the R-package pheatmap ver. 1.0.12 with modified parameters of scaling per row.

### T-DNA regions

The IbT-DNA1 and IbT-DNA2 sequences proposed by Tina Kyndt et al. (2015)^80^were searched against the whole genome with BLAST v2.2.28+. At this time, when the query coverage was over 90%, it was considered a significant hit. Next, when primers obtained from the same reference were mapped to the same region using short BLAST (-task blast-short - word_size 4), the region was finally regarded as a T-DNA region.

## Acknowledgments

We thank the bioinformatics team of PHYZEN (www.phyzen.com) for their technical assistances with the RNAseq analysis. We also thank the Kazusa DNA Research Institute team, Shigemi Sasamoto, Hisano Tsuruoka, Chiharu Minami, Akiko Watanabe, Yoshie Kishida, Mistuyo Kohara, Akiko Komaki, Akiko Obara, Taeko Shibasaki, Rie Aomiya, Takaharu Kimura and Hisako Ichihar for their technical assitances with genome assembly works.

This research was supported by a grant from the National Agricultural Genome Program of Korea (PJ01033901, PJ01033902), the Earmarked Fund for CARS-10-Sweetpotato (CARS-10) in China and Kazusa DNA Research Institute Foundation.

## Data availability

The assembled genome sequences have been submitted to the DDBJ/ENA/NCBI public sequence databases under the BioProject ID PRJDB14817. The assembled genome and gene sequences are available at Plant GARDEN (https://plantgarden.jp/ja/list/t35884/genome/t35884.G002 and https://plantgarden.jp/ja/list/t4120/genome/t4120.G001).

---

**Fig. S1.**
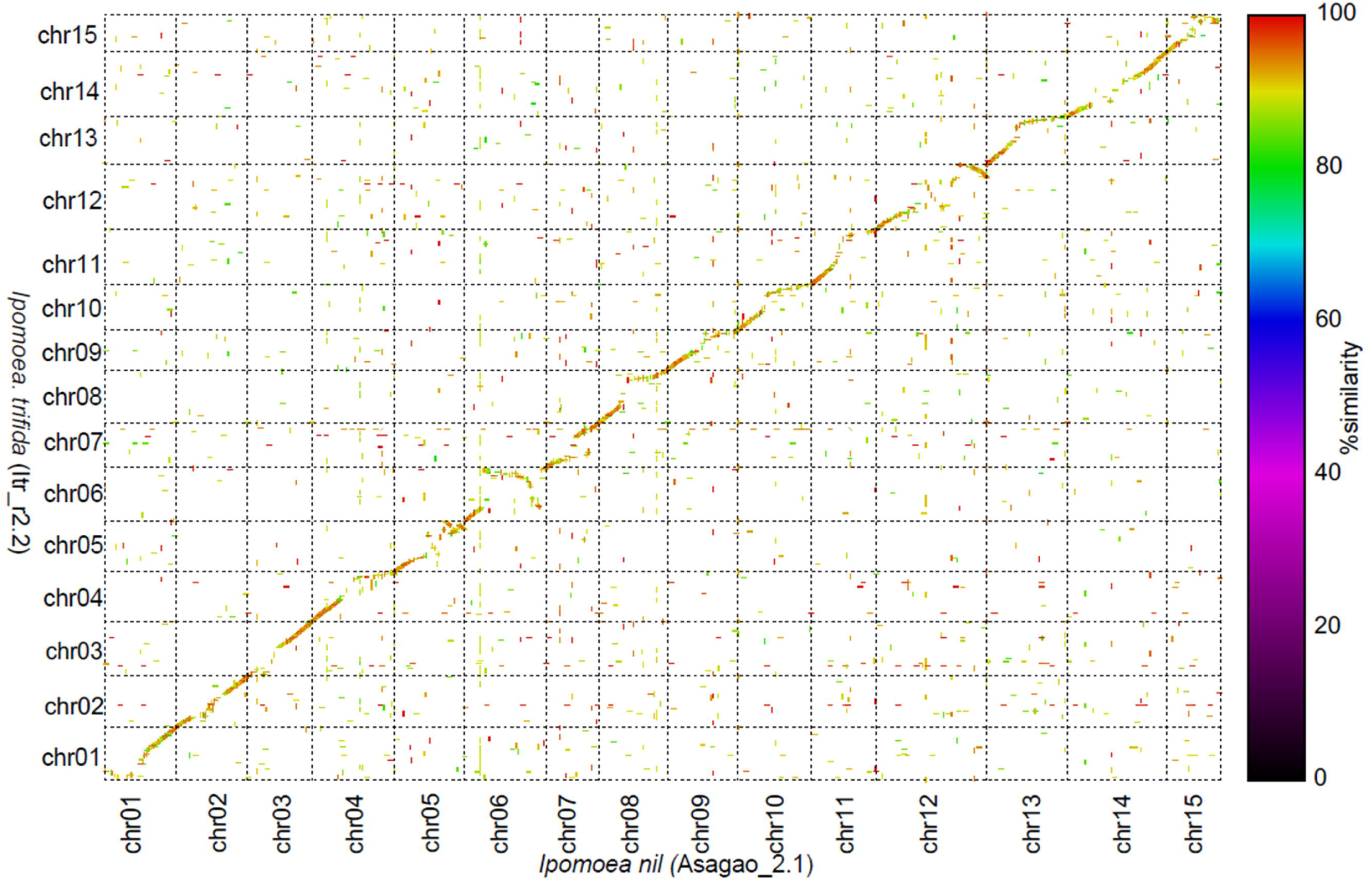
Genome sequence comparison between *I. nil* (Asagao_r2.1) and *I. trifida* (ltr_r2.2).

**Fig. S2.**
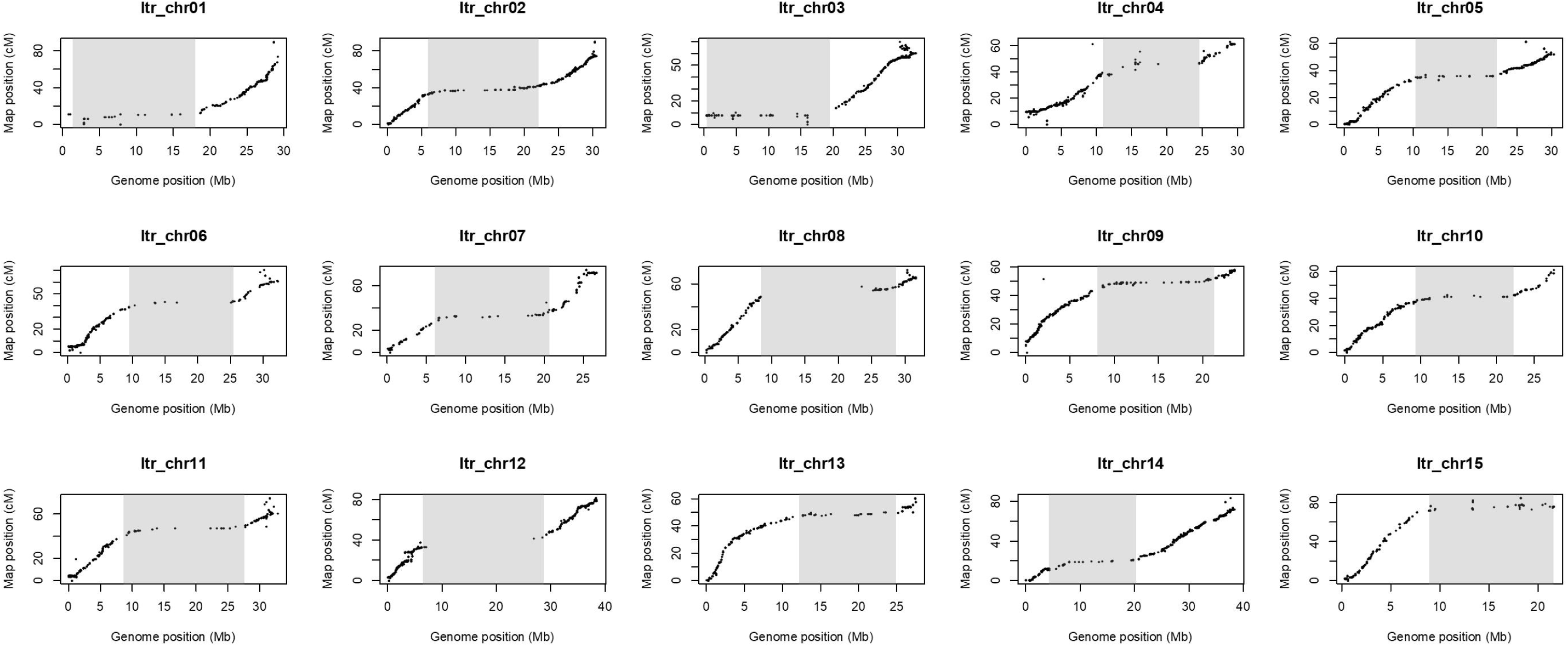
Correlation between physical and genetic distances on the *I. trifida* genome. The genetic distances were calculated from the 0431-1 x Mx23-4 F1 linkage map. Gray boxes represent regions where recombination was suppressed.

**Fig. S3.**
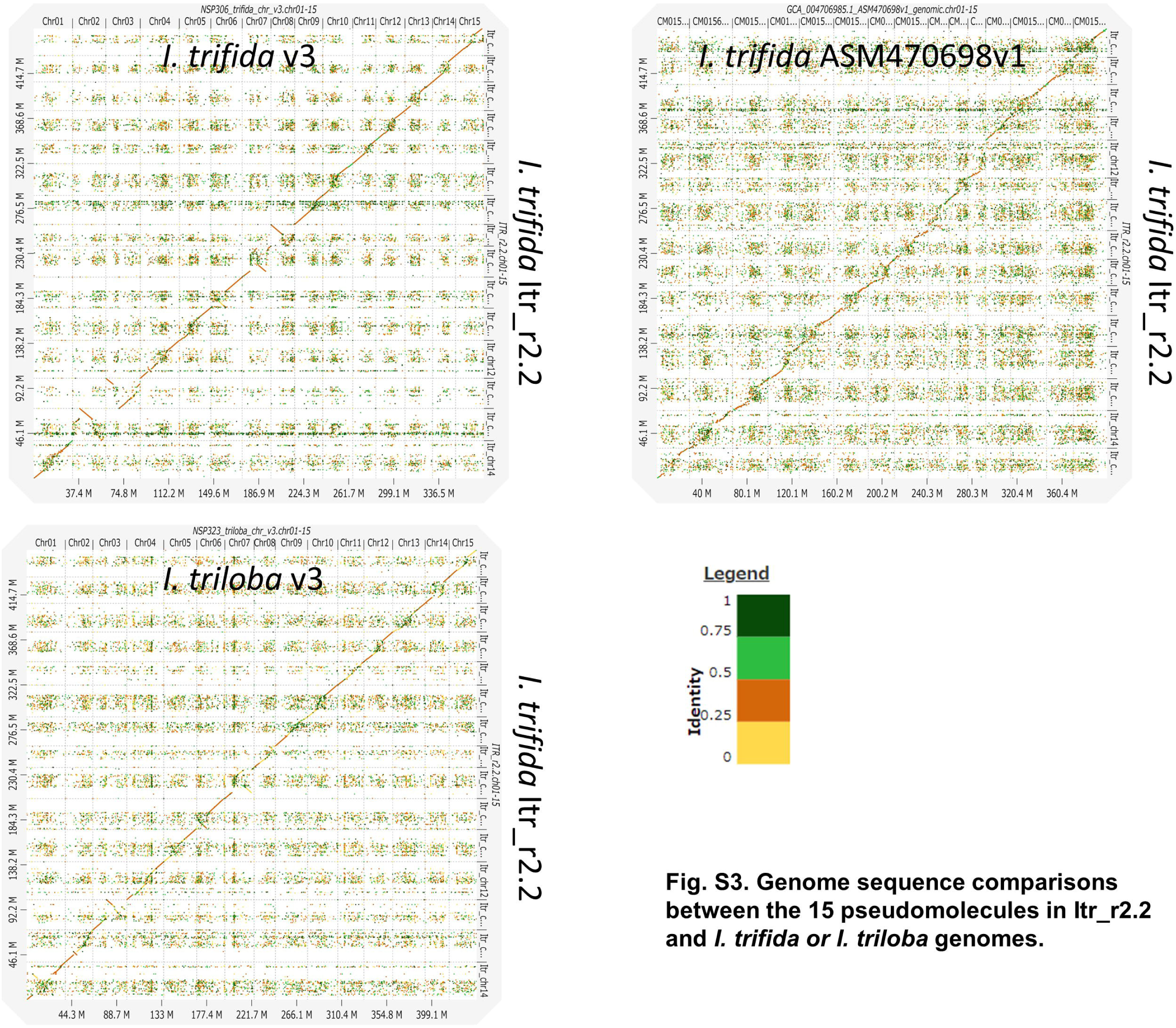
Genome sequence comparisons between the 15 pseudomolecules in ltr_r2.2 and *I. trifida or I. triloba* genomes.

**Fig. S4.**
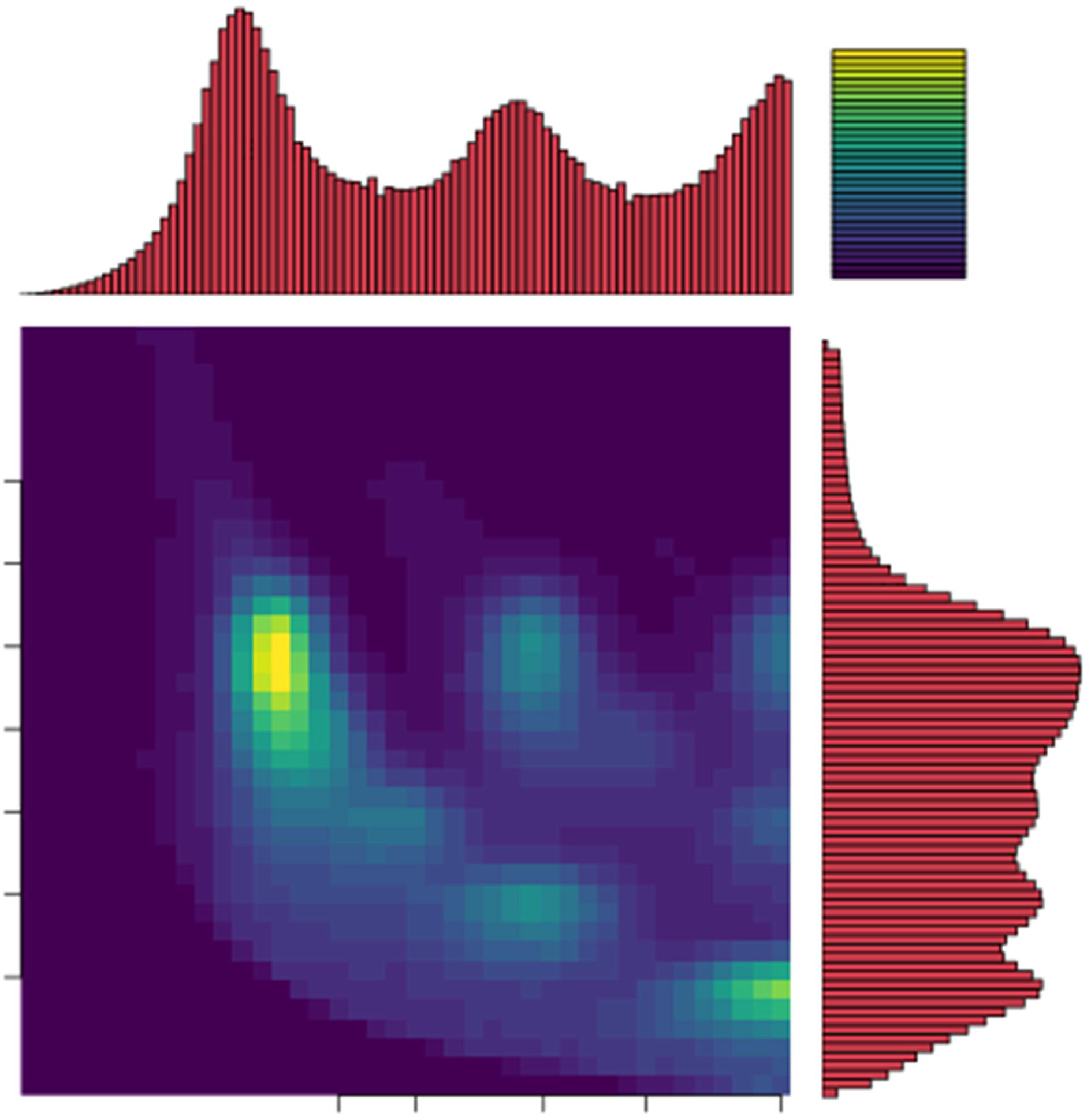
Smudgeplot for *I. batatas* Xushu 18 on the log10 scale. Kmers = 21, 1n = 92. A and B represent homozygous and heterozygous kmers, respectively.

**Fig. S5.**
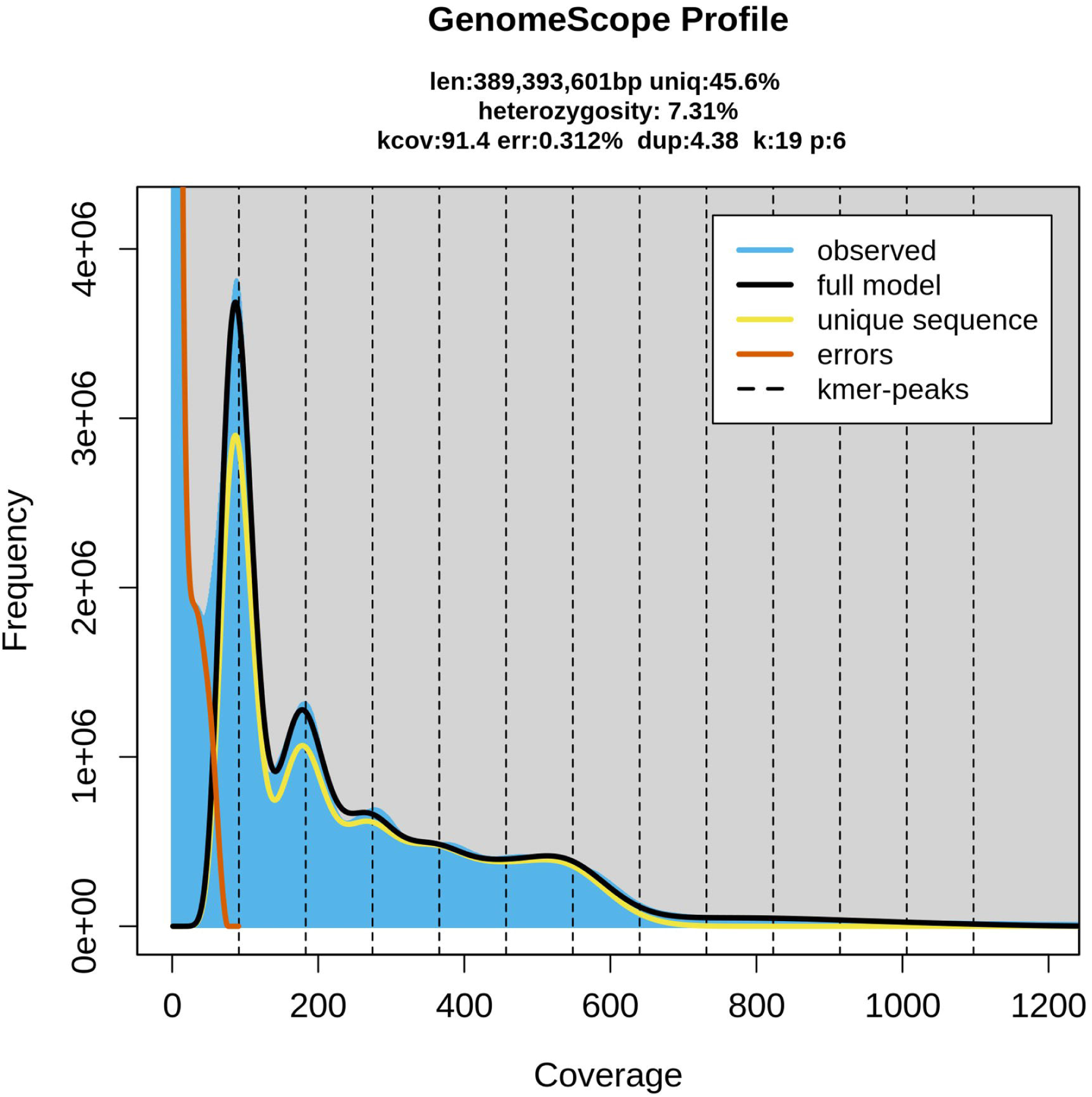
Genome size and heterozygosity estimation using GenomeScope2.0 with the distribution of the number of distinct kmers (kmer = 19, cutoff= 1,000,000) with the given multiplicity values.

**Fig. S6.**
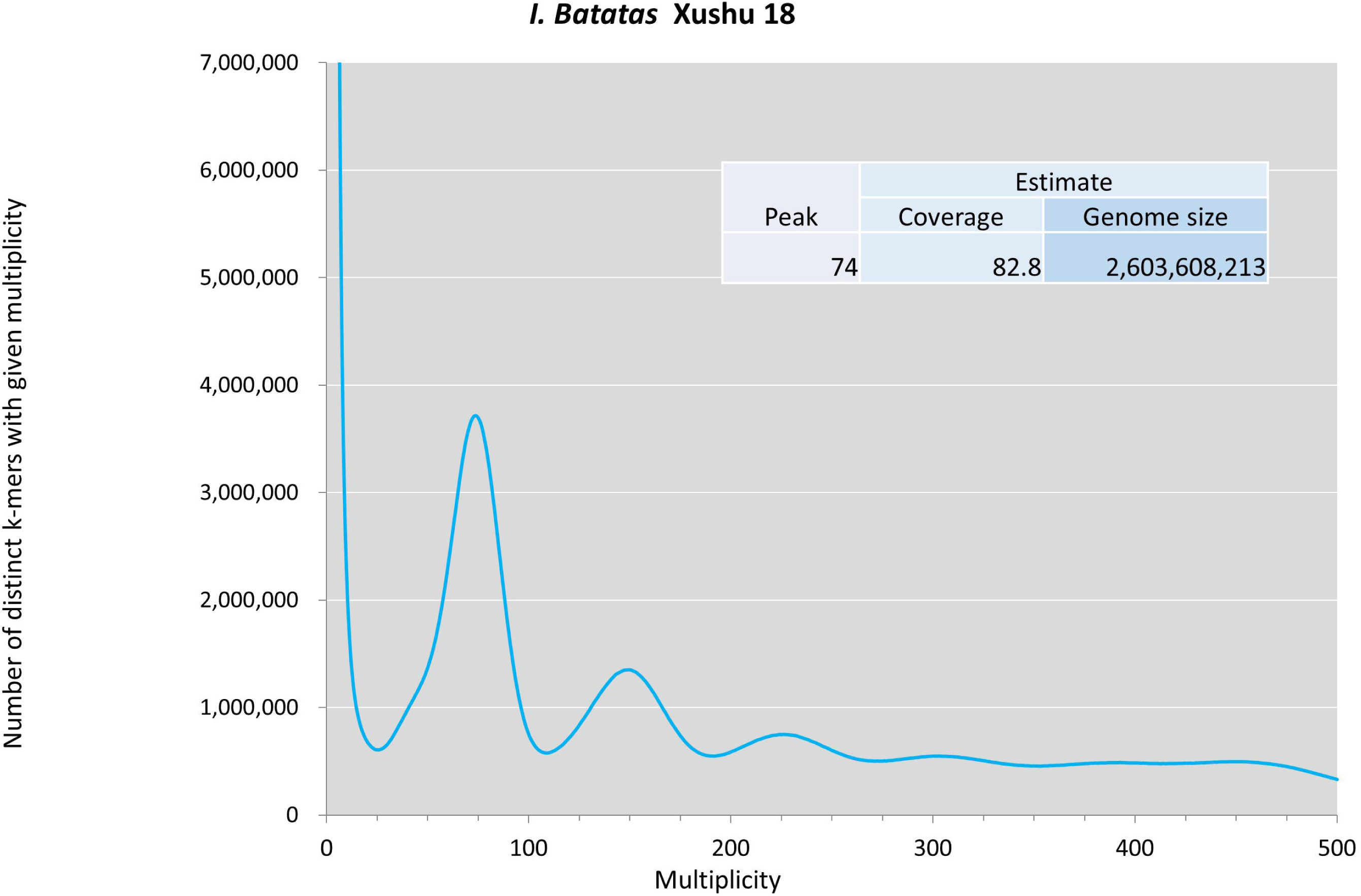
Genome size estimation of *I. batatas* (6X), Xushu 18, using jellyfish with the distribution of the number of distinct kmers (kmer = 17) with the given multiplicity values.

**Fig. S7.**
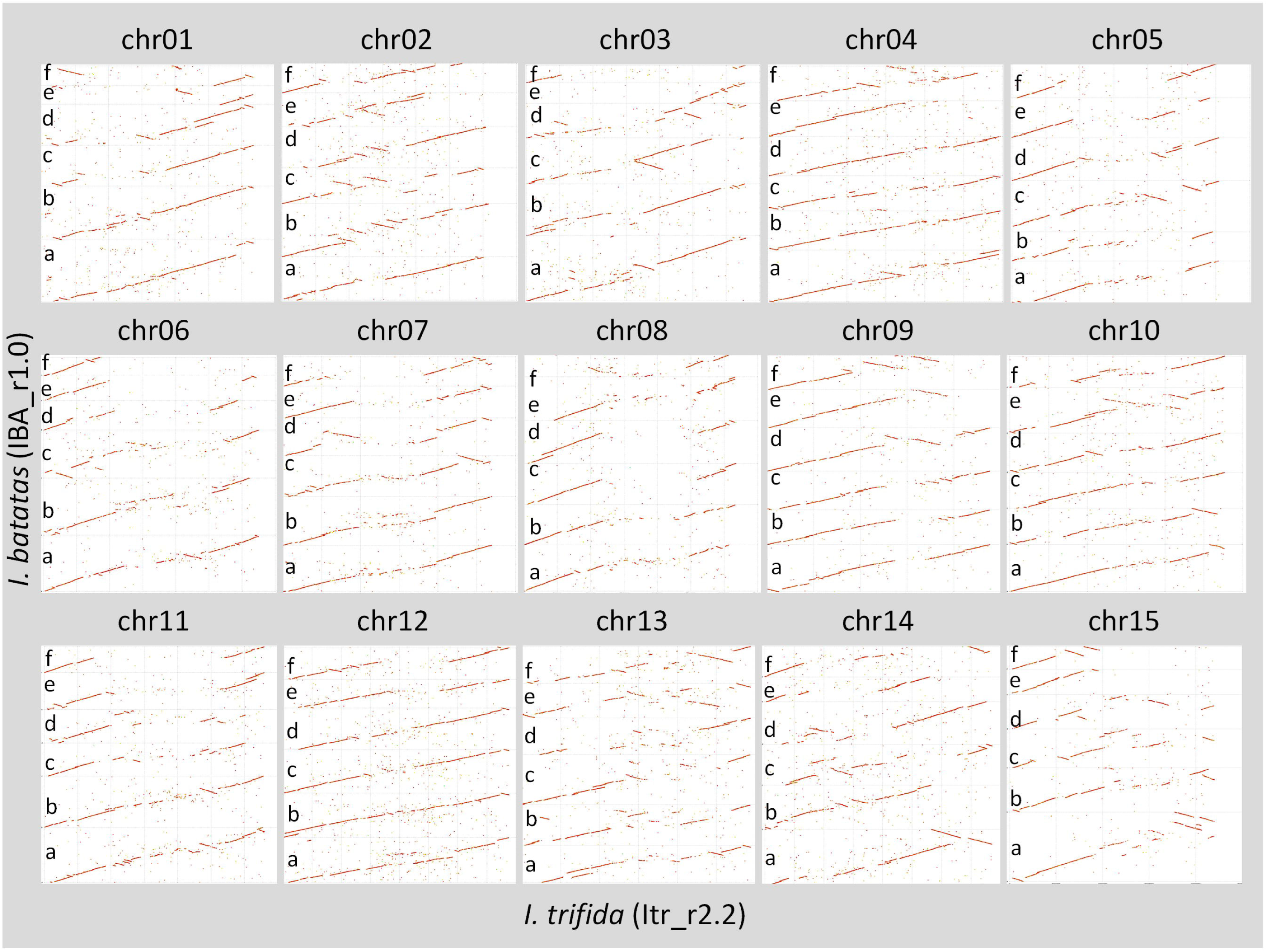
Results of sequence comparison between *I. trifida* (ltr_r2.2) and *I. batatas* (1BA_r1.0) by Nucmer.

**Fig. S8.**
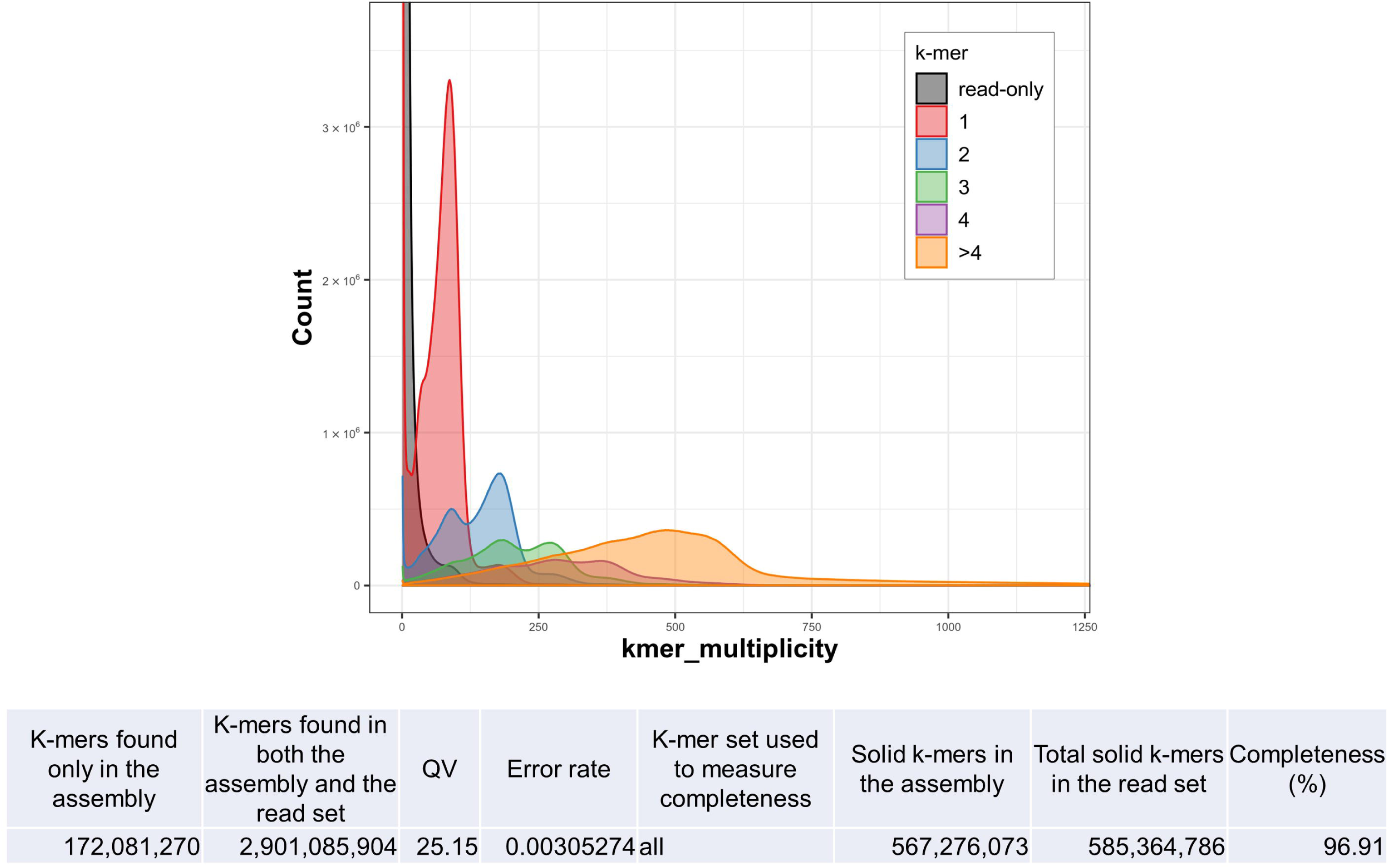
Merqury copy number spectrum plots for the assembled genome, 1BA_r1.0 (including 90 chr and unplaced scaffolds). The red, blue, green, purple, and orange plots represent k-mer multiplicity peaks with x1, x2, x3, x4, and x >4 copy sequences in 1BA_r1.0, respectively. The gray plot represents k-mers identified only on paired-end reads of the sequenced line, Xushu 18. The maximum values of count and k-mer multiplicity calculated by Merqury are 6.69E+09 and 1.13E+08, respectively. The figure illustrates partial data in order to show the distribution clearly.

**Fig. S9.**
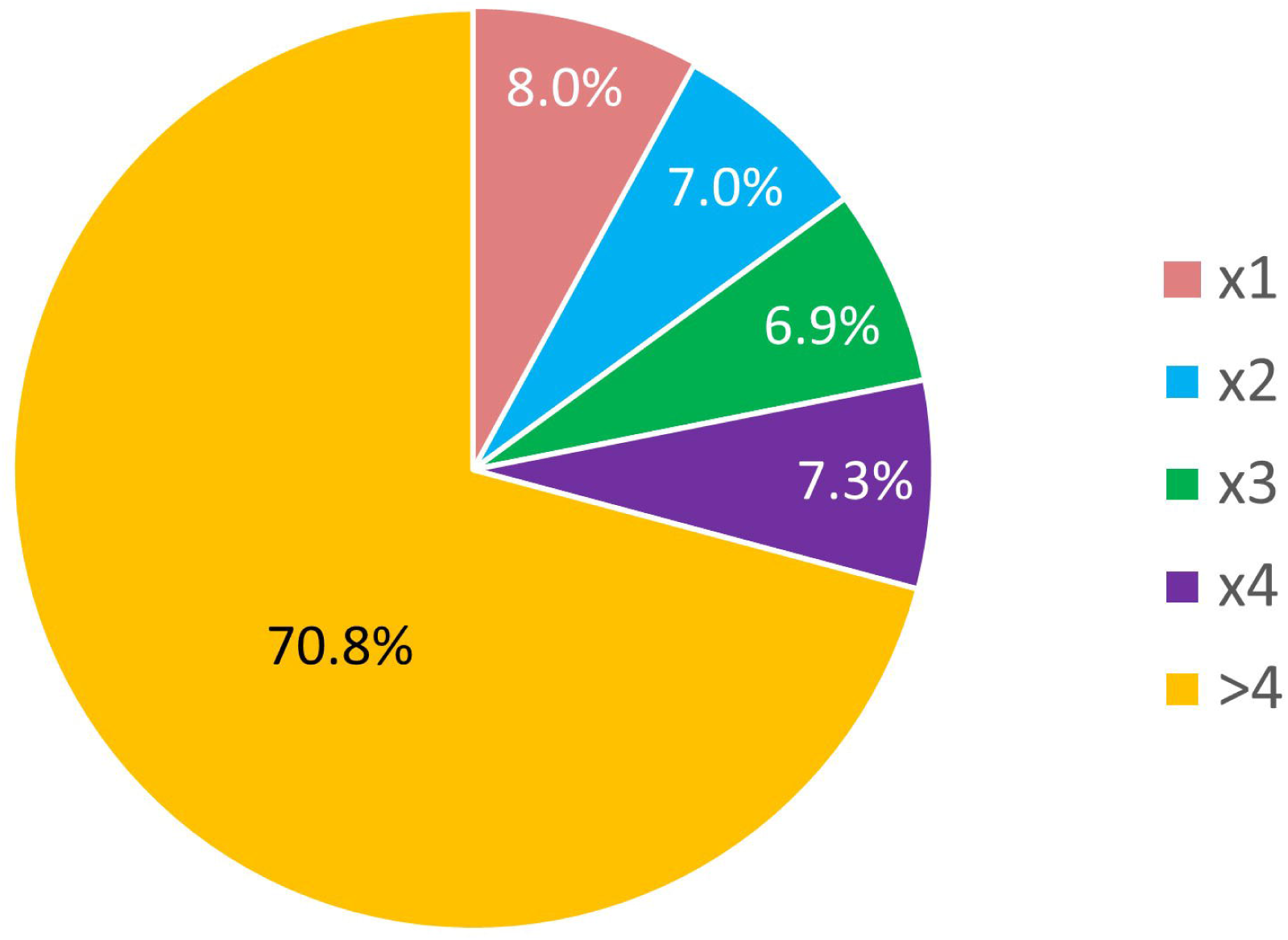
Ratios of genome sequence reads (DRX****) classified as x1, x2, x3, x4, and x >4 copy by Merqury. The ratios were calculated by comparing the multiplication of k-mer multiplicities and counts.

**Fig. S10.**
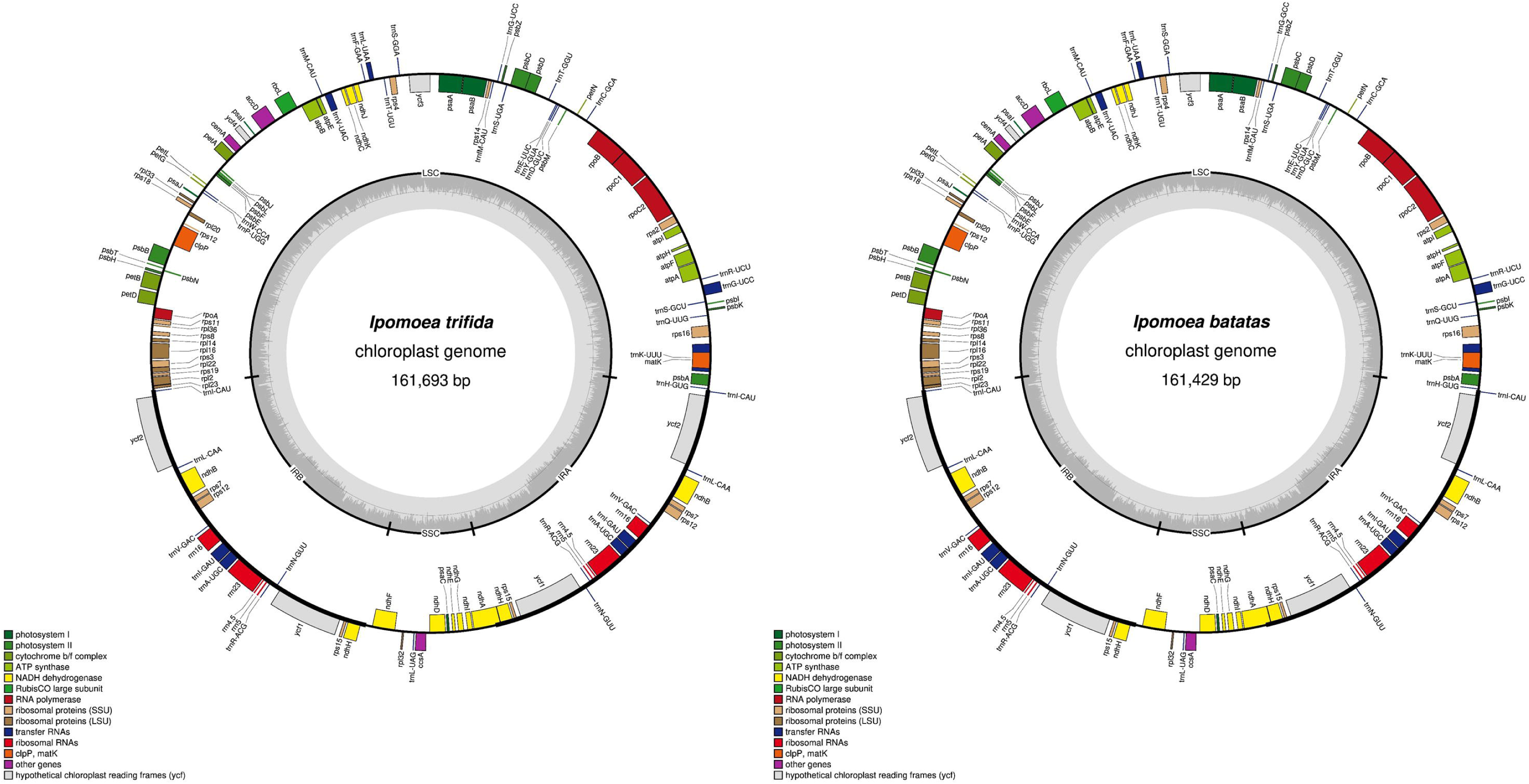
Chloroplast genome map of *I. trifida* Mx23Hm (left) and *I. batatas* Xushu 18 (right). The outer circle shows the position of genes. The inner circle represents GC content.

**Fig. S11.**
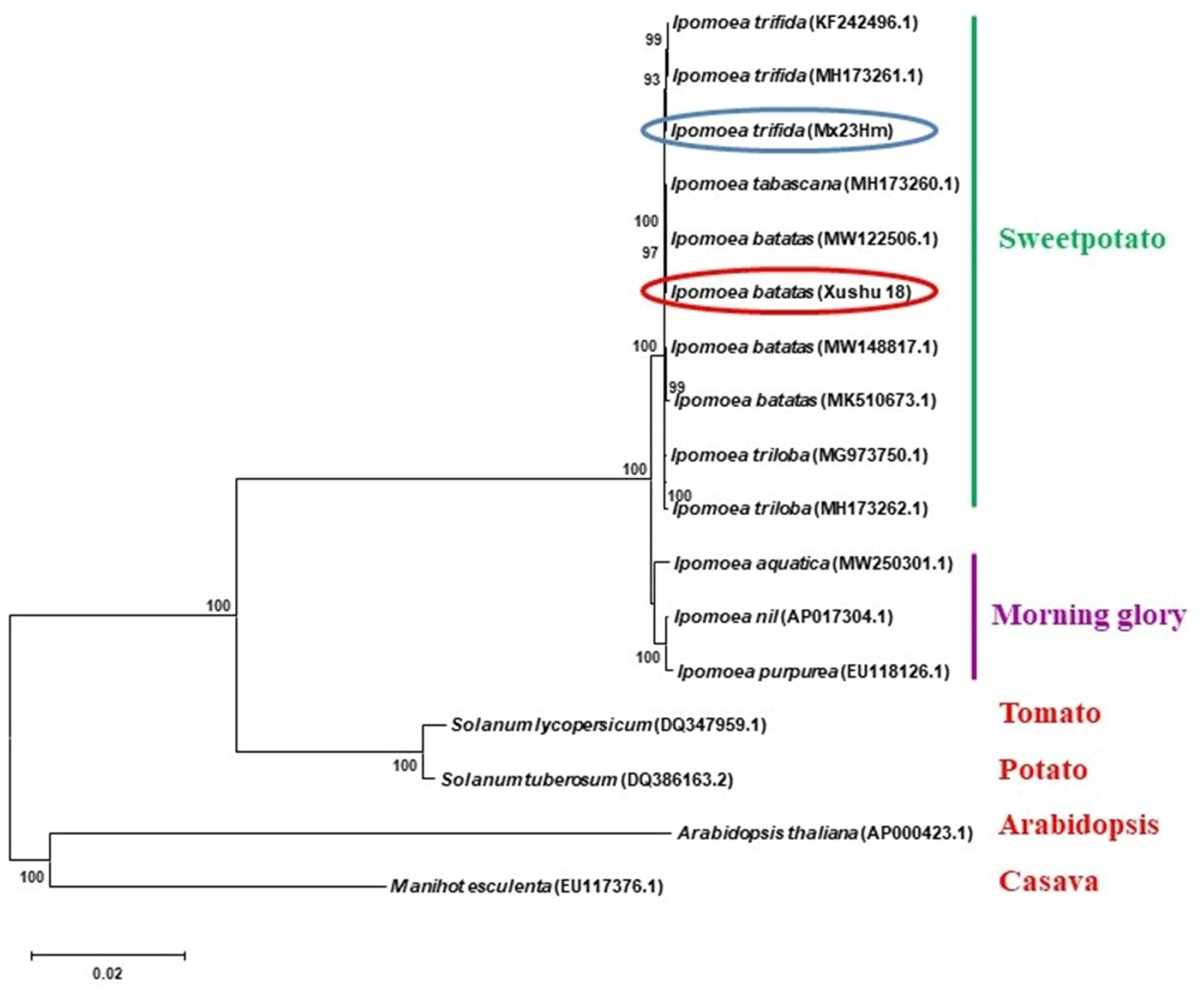
Phylogenetic analysis of chloroplast genome. Phylogenetic analysis of 17 species using 68 protein-coding sequences of the chloroplast genome. Outgroups were *Manihot esculenta* and *Arabidopsis thaliana.* A phylogenetic tree was generated using the maximum-likelihood method and 1000 bootstrap replicates by MEGA11. Genome list: *lpomoea. batatas* Xushu 18, *I. trifida* Mx23Hm, *I. batatas* MW148817, *I. batatas* MW122506, *I. batatas* MK510673, *I. trifida* MH173261, *I. trifida* KF242496, *I. nil* AP017304, *I. triloba* MH173262, *I. triloba* MG973750, *I. tabascana* MH173260, *I. purpurea* EU118126, *I. aquatica* MW250301, *Solanum lycopersicum* DQ347959, S. *tuberosum* DQ386163, *Manihot esculenta* EU117376, *Arabidopsis thaliana* AP000423

**Fig. S12.**
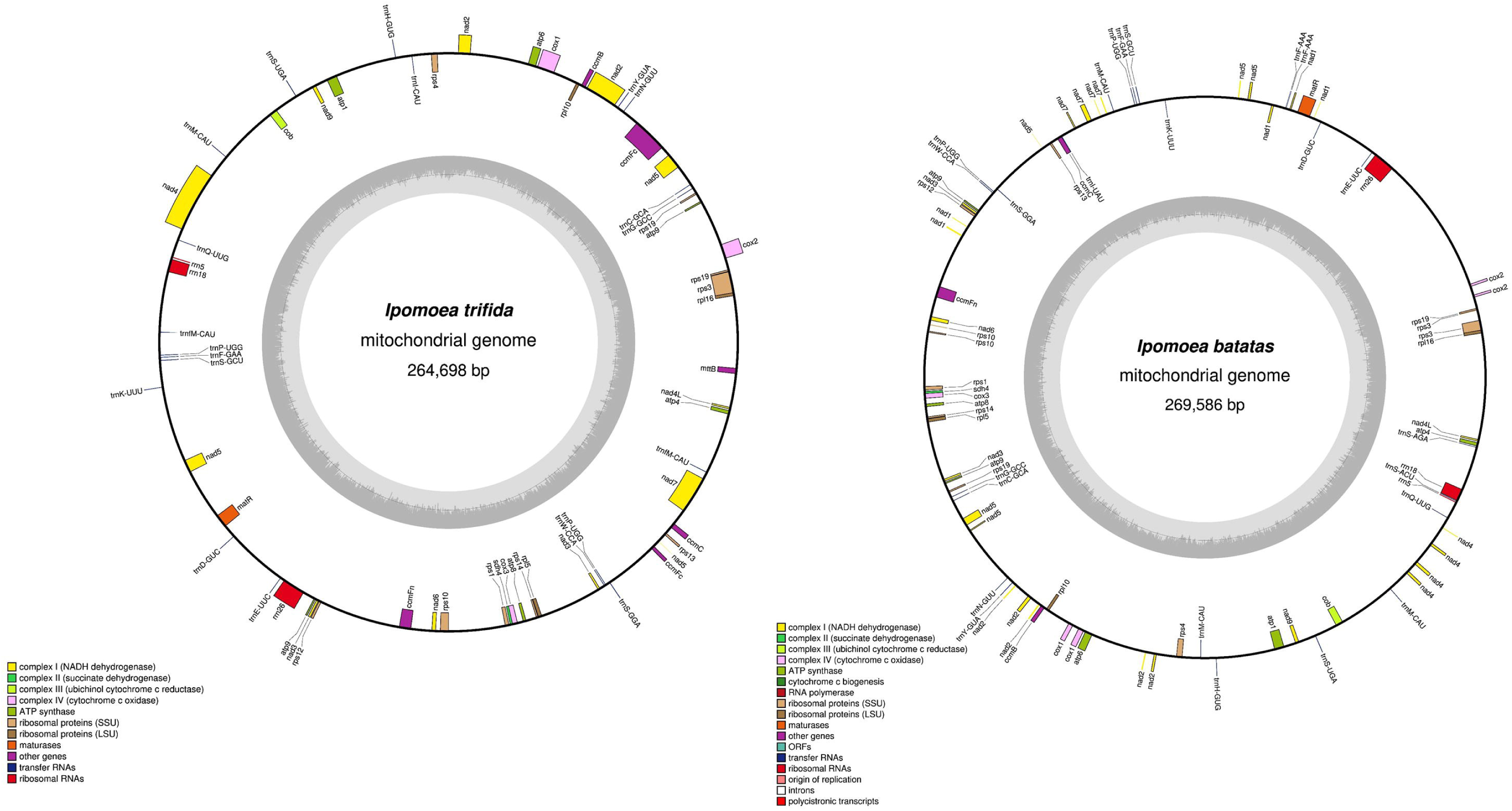
Mitochondrial genome map of *I. trifida* Mx23Hm (left) and *I. batatas* Xushu 18 (right). The outer circle shows the gene positions. The inner circle represents GC content.

**Fig. S13.**
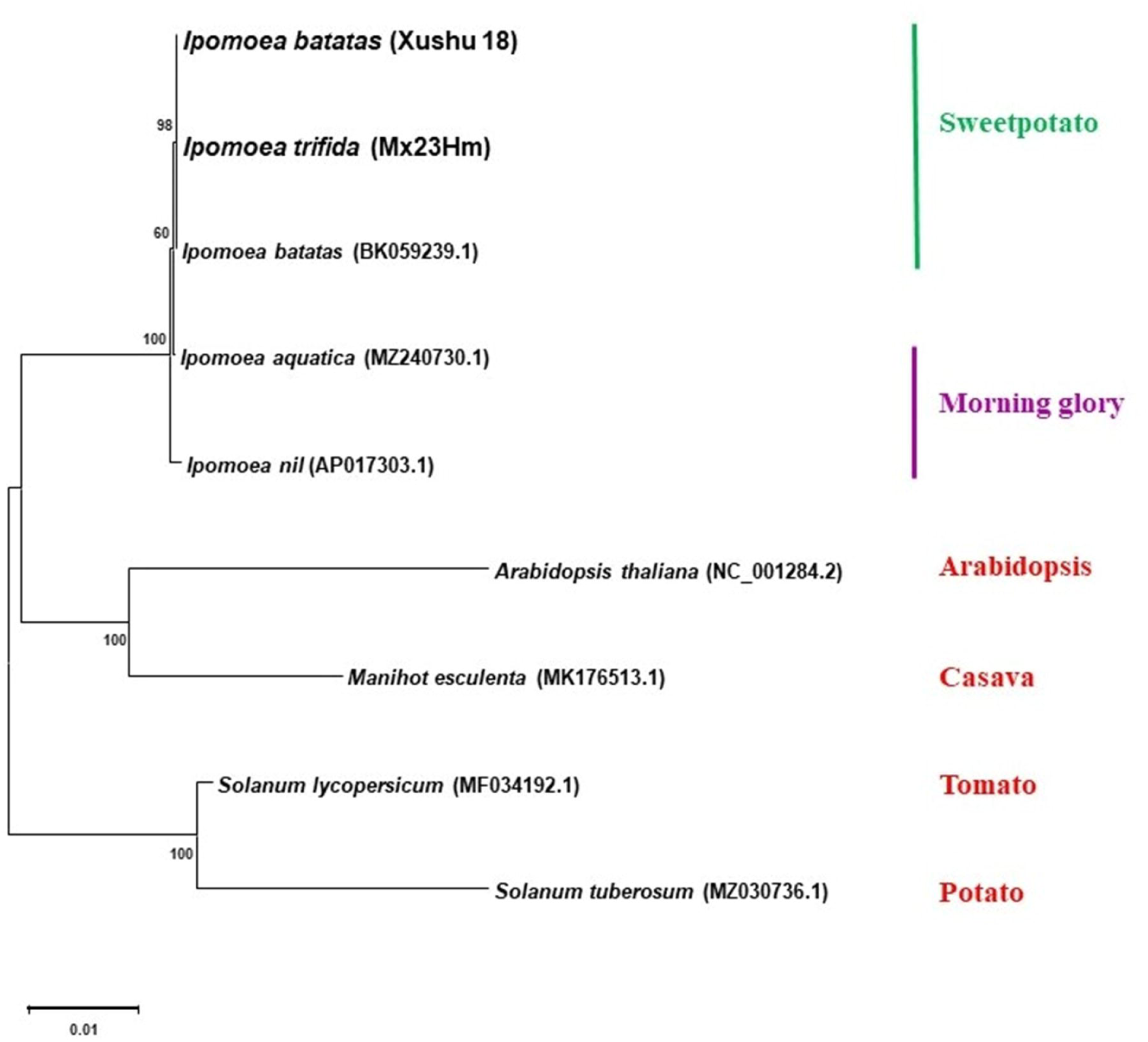
Phylogenetic analysis of mitochondrial genome. Phylogenetic analysis of 9 species using 10 protein-coding sequences of the mitochondrial genome. A phylogenetic tree was generated using the maximum-likelihood method and 1000 bootstrap replicates by MEGA11. Genome list: *lpomoea batatas* Xushu 18, *I. trifida* Mx23Hm, *I. batatas* BK059239, *I. nil* AP017303, *I. aquatica* MZ240730, *Solanum lycopersicum* MF034192, *S. tuberosum* MZ030736, *Manihot esculenta* MK176513, *Arabidopsis thaliana* NC_001284.

**Fig. S14.**
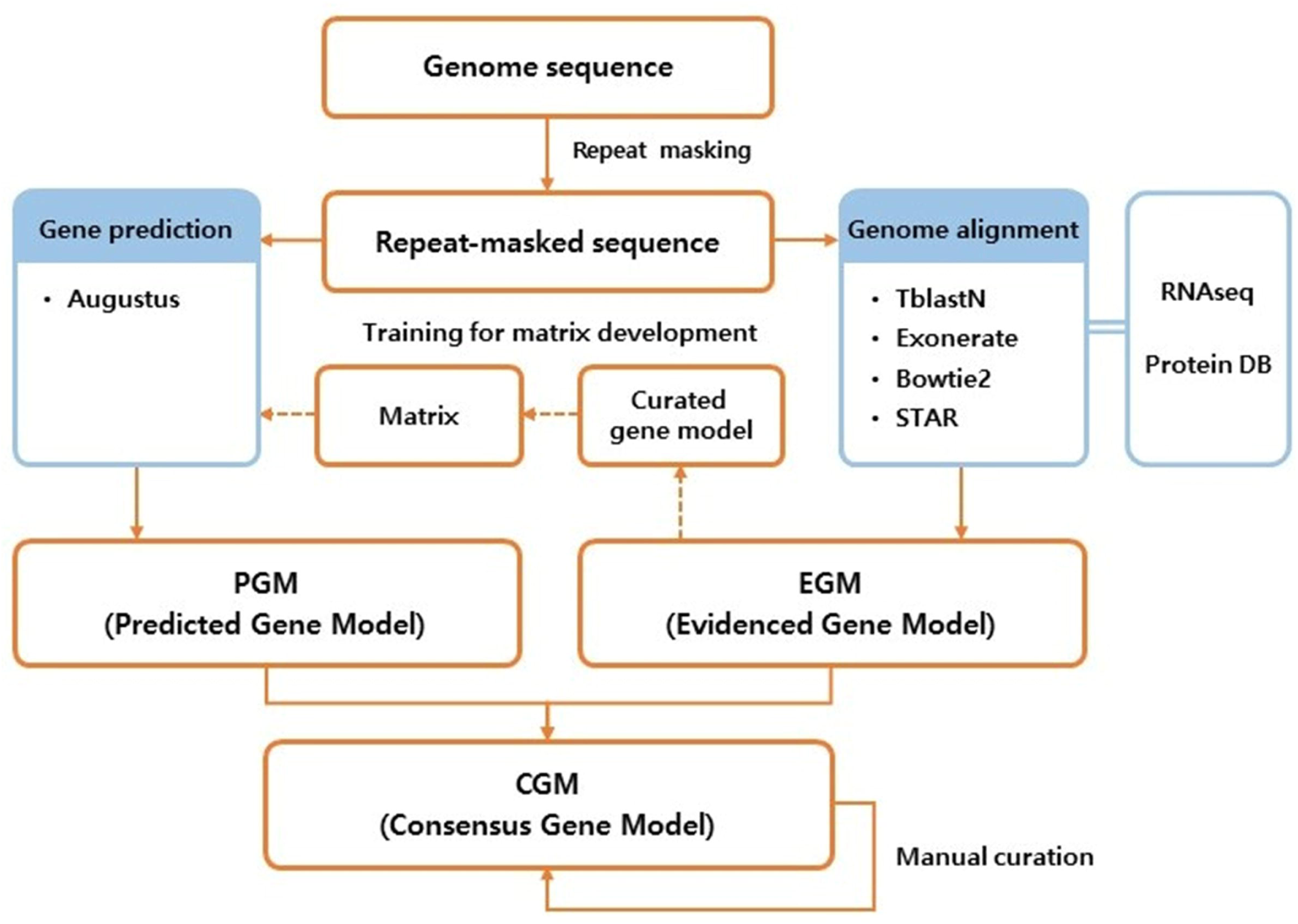
Schematic workflow for the gene prediction of *I. trifida* (ltr_r2.2) and *I. batatas (*IBA_r1.0).

**Fig. S15.**
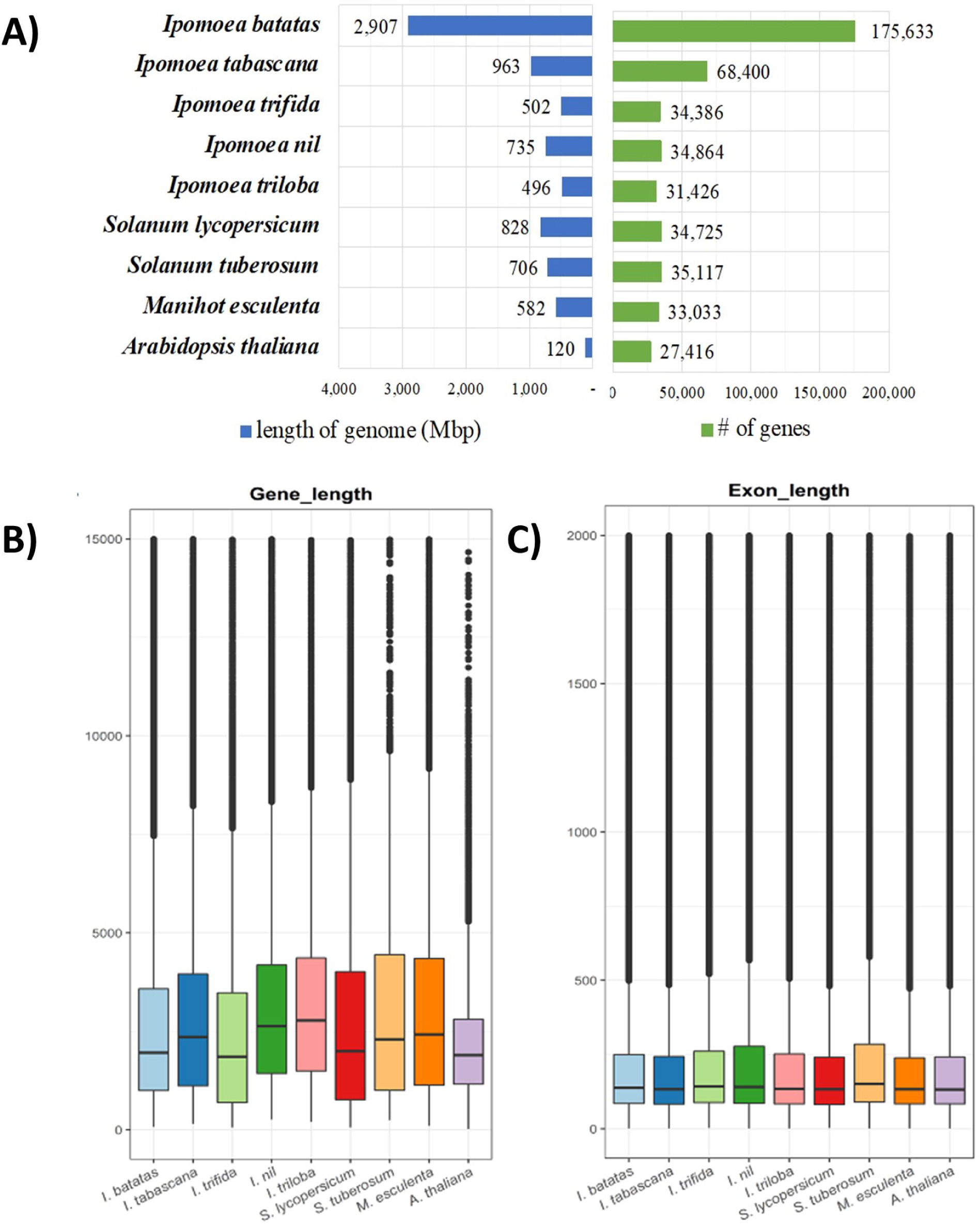
The assembled genome length and protein coding gene numbers in *I. batatas* (IBA_r1.0), *I. trifida* (ltr_r2.2), and 7 other species. The genome versions except *I. batatas* and *I. trifida* are listed in Table S13. **A**) Genome assembly lengths and numbers of predicted genes. **B**) Exon lengths of protein coding genes. **C**) Exon lengths.

**Fig. S16.**
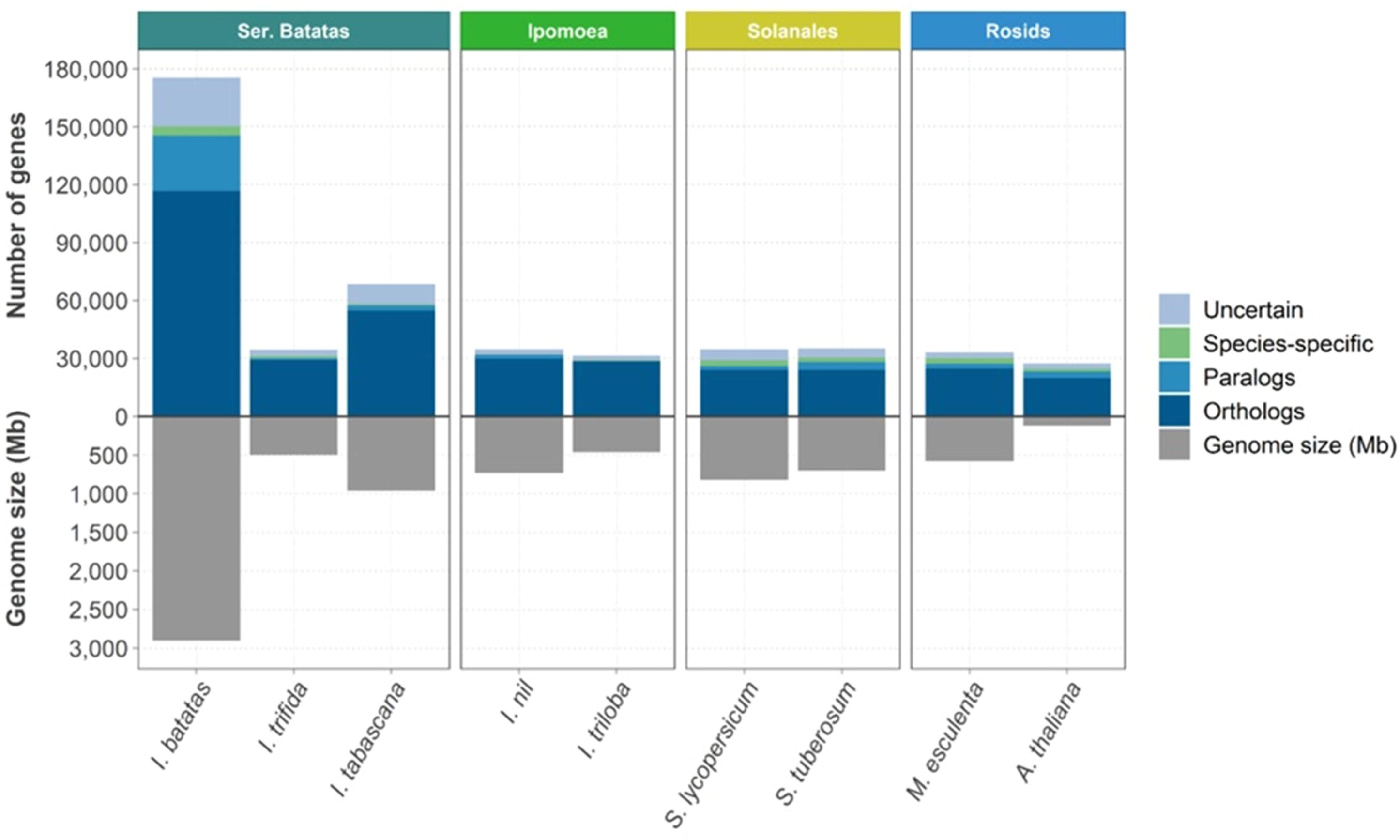
The results of orthologous analysis among the 9 genomes. Ortholog analysis based on protein sequence homology in the 9 genomes. The lower tier represents genome size and the upper tier represents the number of genes. Orthologous genes: orthologous genes among 9 species; specific: genes only in their species; else: genes that have sequence similarity to other species but are not established due to ortholog relationships.

**Fig. S17.**
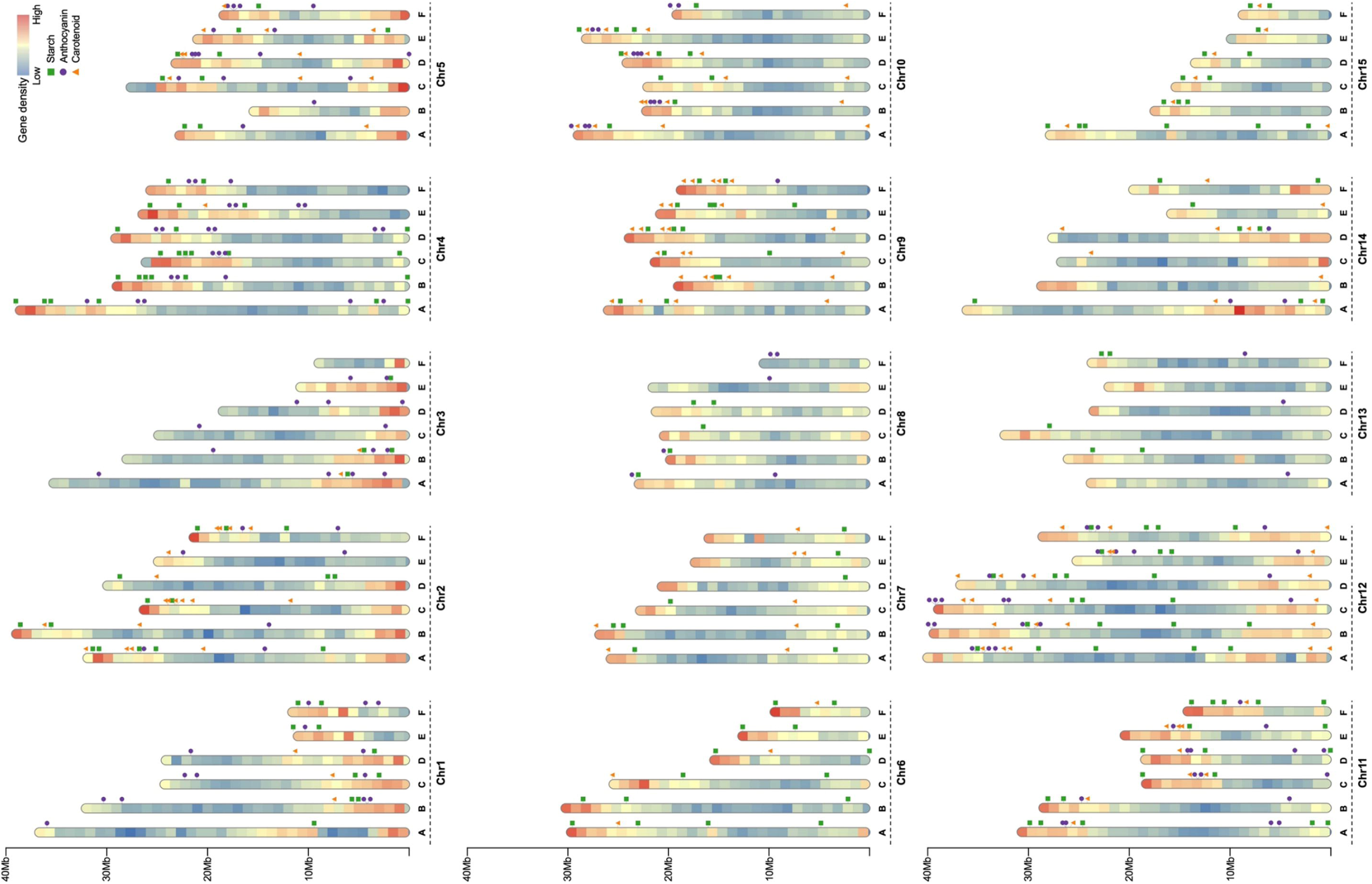
Gene map of stratch synthesis genes, anthocyanin synthesis genes, and carotenoid synthesis genes in *I. batatas*.

**Fig. S18.**
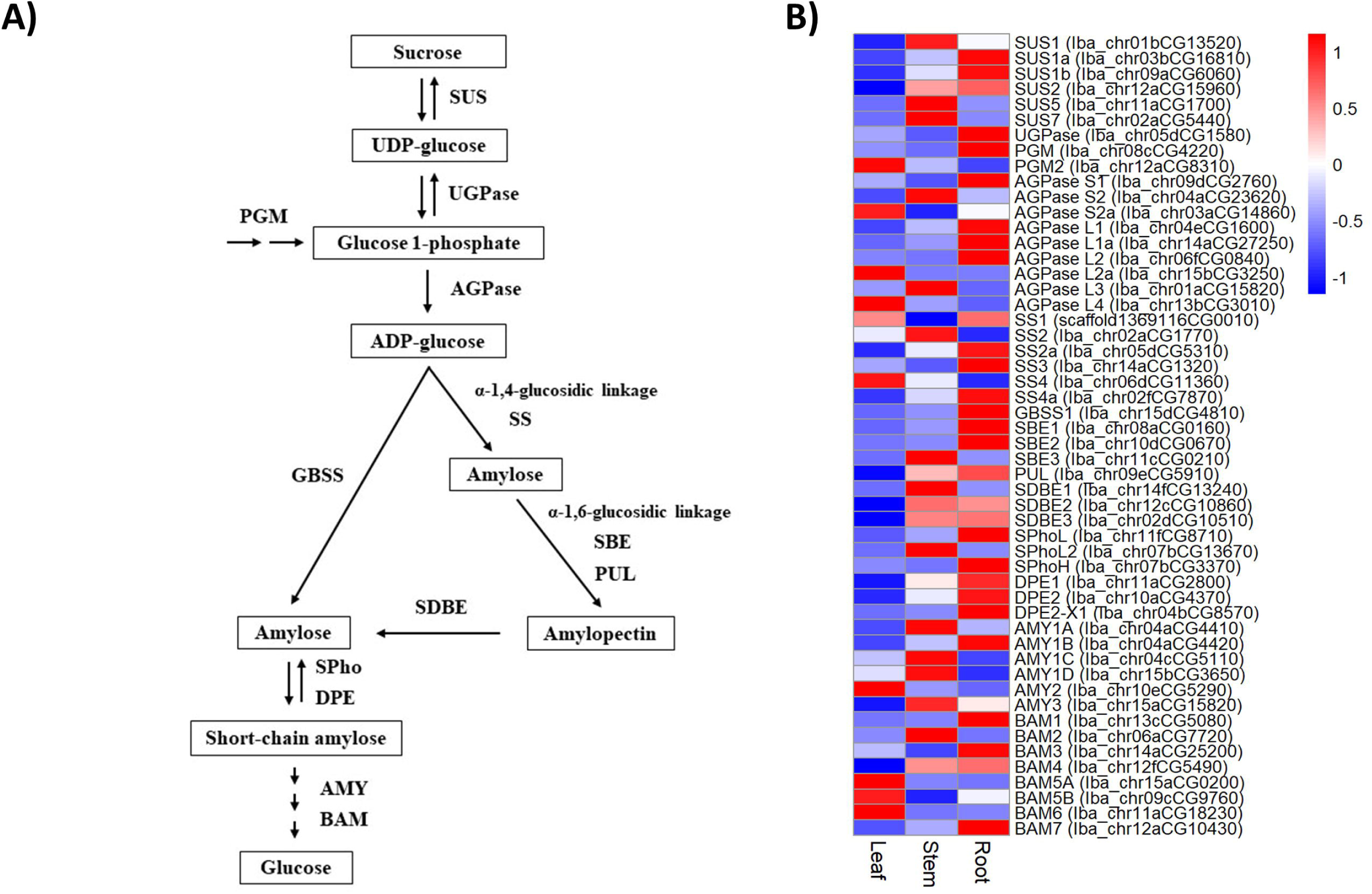
Schematic presentation of the starch synthesis pathway in sweetpotato and expression patterns of starch pathway genes. **(A)** Starch synthesis pathway in sweetpotato. SUS, sucrose synthase; UGPase, UDP-glucose pyrophosphorylase; PGM, phosphoglucomutase; AGPase, ADP-glucose pyrophosphorylase ; SS, starch synthase; GBSS, granule-bound starch synthase; SBE, starch branching enzyme; PUL, pullulanase; SOBE, starch debranching enzyme; SPho, starch phosphorylase; OPE, a-1,4-glucanotransferase; AMY, a-amylase ; BAM, ­ amylase. **(B)** Expression heatmap of the starch synthesis pathway genes in 3 tissues. A heatmap was generated with the average FPKM values in 2 replicates of genes using the R-package pheatmap ver. 1.0.12 with modified parameters of scaling per row to visualize gene expression peaks among the different samples.

**Fig. S19.**
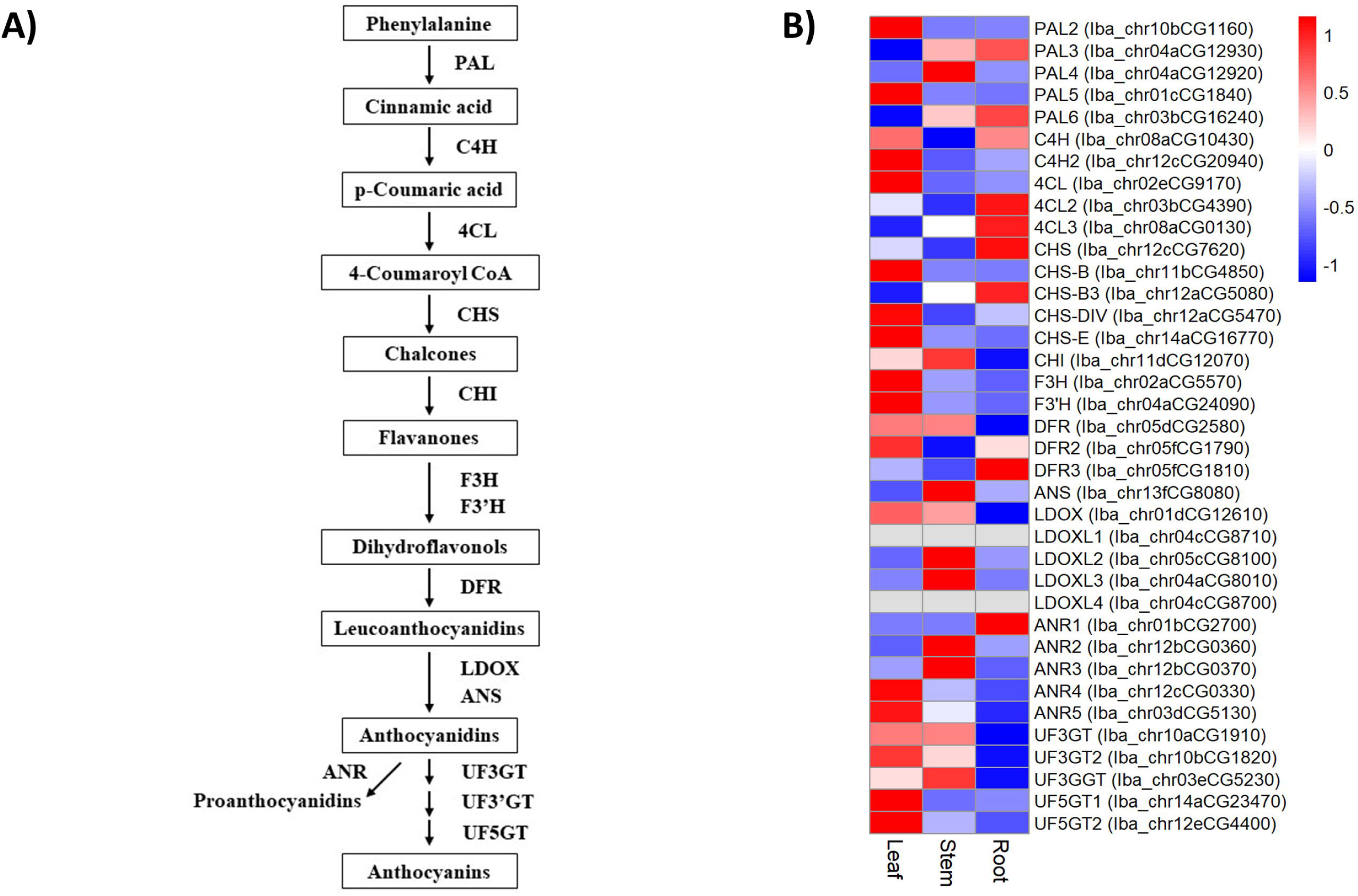
Schematic presentation of the anthocyanin synthesis pathway in sweetpotato and expression patterns of the starch synthesis pathway genes. **(A)** Anthocyanin synthesis pathway in sweetpotato. PAL, phenylalanine ammonia lyase; C4H, cinnamic acid 4-hydroxylase; 4CL, 4-coumarate CoA ligase; CHS, chalcone synthase; CHI, chalcone isomerase; F3H, flavanone 3-hydroxylase; F3’H, flavonoid 3’-hydroxylase; DFR, dihydroflavonol 4-reductase; LDOX, leucoanthocyanidin dioxygenase; ANS, anthocyanidin synthase; ANR, anthocyanidin reductase; UF3GT, anthocyanidin 3-0 -glucosyltransferase; UF3GGT, anthocyanidin 3-0-glucoside 2’-0 - glucosyltransferase; UF5GT, anthocyanidin 5-0-glucosyltransferase. (B) Expression heatmap of the anthocyanin synthesis pathway genes in 3 tissues.

**Fig. S20.**
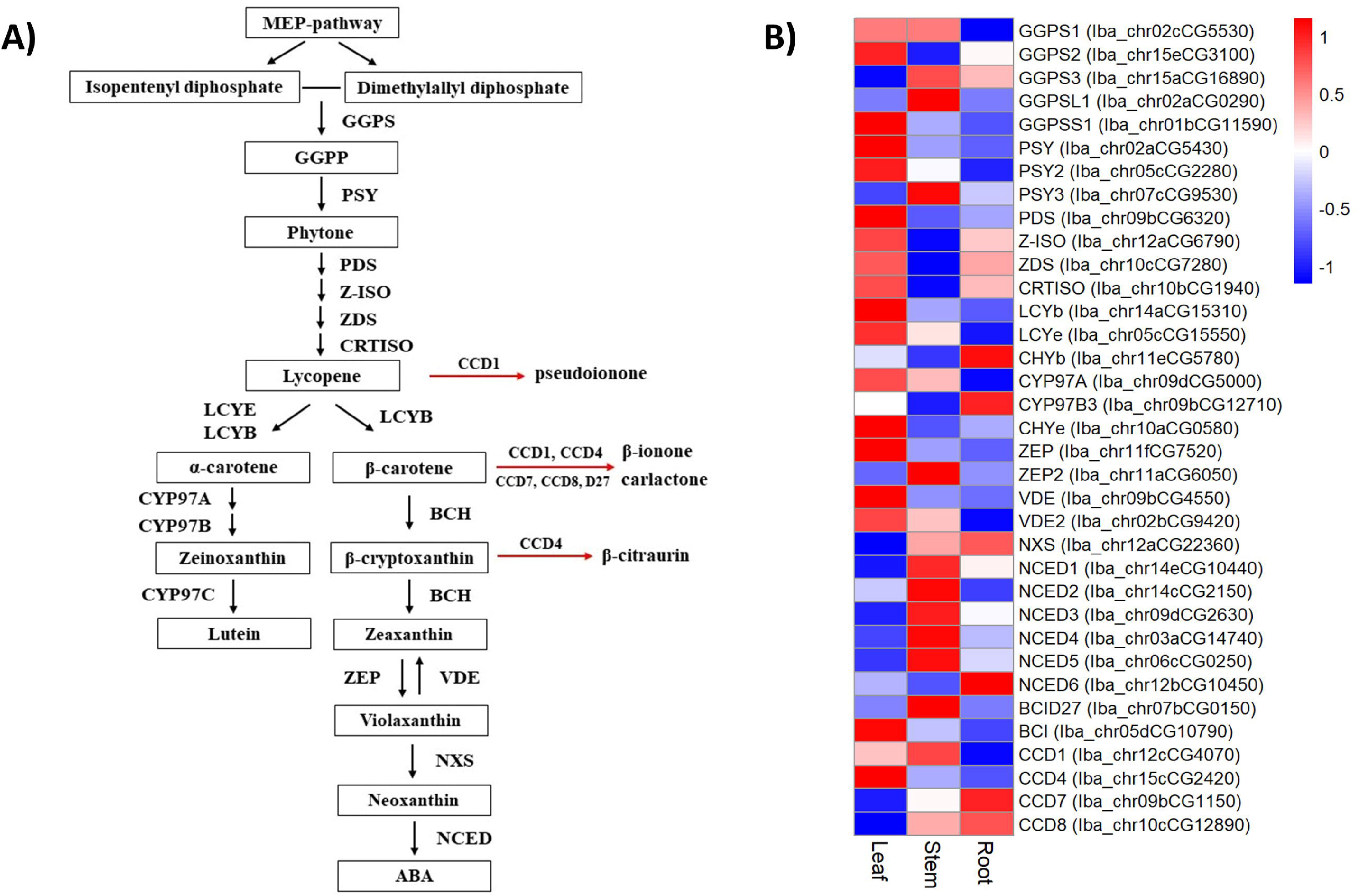
Schematic presentation of the carotenoid synthesis pathway in sweetpotato and expression patterns of the carotenoid synthesis pathway genes. **(A)** Carotenoid synthesis pathway in sweetpotato. MEP, methylerythritol 4-phosphate; IPP, isopentenyl diphosphate; DMAPP, Dimethylallyl diphosphate; GGPP, geranylgeranyl diphosphate; GGPS, geranylgeranyl pyrophosphate synthase; PSY, phytoene synthase; PDS, phytoene desaturase; Z-ISO, zeta carotene isomerase; ZDS, zeta carotene desaturase; CRTISO, carotenoid isomerase; LCYE, lycopene epsilon cyclase; LCYB, lycopene beta cyclase; CYP97A, cytochrome P450 97A; CYP97B, cytochrome P450 97B; CYP97C, cytochrome P450 97C; BCH, beta carotene hydroxylase; ZEP, zeaxanthin epoxidase; VDE, violaxanthin de-epoxidase; NXS, neoxanthin synthase; NCED, 9-cis-epoxycarotenoid dioxygenase; CCD, carotenoid cleavage dioxygenase; BCID27, beta carotene isomerase D27. (B) Expression heatmap of the carotenoid synthesis pathway genes in 3 tissues.

